# Oligodendrocytes use postsynaptic proteins to coordinate myelin formation on axons of distinct neurotransmitter classes

**DOI:** 10.1101/2024.10.02.616365

**Authors:** Natalie J. Carey, Caleb A. Doll, Bruce Appel

**Affiliations:** Section of Developmental Biology, Department of Pediatrics, University of Colorado, Anschutz Medical Campus, Aurora, Colorado, USA, 80445

## Abstract

Axon myelination can tune neuronal circuits through placement and modulation of different patterns of myelin sheaths on distinct types of axons. How myelin formation is coordinated on distinct axon classes remains largely unknown. Recent work indicates neuronal activity and vesicle release promote myelin formation, and myelin-producing oligodendrocytes express canonical postsynaptic factors that potentially facilitate oligodendrocyte-axon interaction for myelin ensheathment. Here, we examined whether the inhibitory postsynaptic scaffold protein Gephyrin (Gphn) mediates selective myelination of specific axon classes in the larval zebrafish. Consistent with this possibility, Gphn was enriched in myelin on GABAergic and glycinergic axons. Strikingly, in *gphnb* deficient larvae, myelin sheaths were longer specifically on GABAergic axons, and the frequency of myelin placement shifted toward glutamatergic axons at the expense of GABAergic axons. Collectively, our results indicate that oligodendrocytes use postsynaptic machinery to coordinate myelin formation in an axon identity-dependent manner.

## INTRODUCTION

Oligodendrocytes (OLs) are glial cells in the central nervous system (CNS) that produce myelin, a lipid-rich membrane that wraps around axons to provide metabolic and trophic support and increase action potential velocity. A single OL can myelinate dozens of axons simultaneously^1^, including different classes of axons defined by distinct neurotransmission profiles. Because different neuron types have different axon lengths, firing rates, and energetic demands, the amount and composition of myelin on axons can play a crucial role in signal timing within a neural circuit.

Neural circuits require a balance of excitatory and inhibitory influence to achieve regulated output, such as the coordinated locomotion generated by the spinal cord. Canonically, glutamatergic neurons provide excitatory input and γ-amino butyric (GABAergic)/glycinergic neurons provide inhibitory influence on circuit output^2–4^. Critically, glutamatergic, GABAergic, and glycinergic neurons signal through unique molecular machinery where their axon terminals create synapses with the appropriate postsynaptic terminal. Postsynaptic scaffold proteins provide specificity for synapse formation by anchoring receptors and cell adhesion molecules that are enriched at unique synapses. Postsynaptic Density 95 protein (PSD95) is the primary scaffold protein at excitatory glutamatergic synapses, whereas Gephyrin (Gphn) is the postsynaptic scaffold at inhibitory GABAergic and glycinergic synapses^3–9^. This specificity of synaptic communication is necessary in complex circuits to coordinate neuronal firing and generate functional behaviors such as locomotion.

Remarkably, OLs produce myelin sheaths with variable lengths and thicknesses on individual axons^10–11^, and myelin patterns on distinct classes of axons vary across neuron type and brain region^12–16^. What mechanisms might convey specificity in myelin formation on distinct axon classes? One possibility is that OLs engage with axons using mechanisms similar to synapse formation where a myelin sheath contacts an axon at an axo-sheath interface. Several findings support this possibility. First, neuronal activity promotes myelin formation through vesicle release along the axon^15,17–19^. This vesicular release is accompanied by axonal Ca^2+^ events at sites where myelin growth will subsequently occur^15^. Second, gene expression profiling studies show that OL lineage cells (OLCs) express many genes that encode postsynaptic proteins such as PSD95 and Gphn^20–23^. Third, interfering with postsynaptic protein function in OLs disrupts myelin formation and maintenance^24–25^. And fourth, an OL precursor cell (OPC)-specific knockout of GABA_A_R γ2 altered myelin profiles on fast-spiking, GABAergic PV interneurons and their subsequent firing rate without impacting the myelin or firing rate of neighboring, glutamatergic spiny stellate cell interneurons^26^. Therefore, OLs and their myelin sheaths are uniquely positioned to modulate neural circuit output by regulating signal timing and consequently alter the strength or frequency of circuit signals or eliminate output altogether^27–29^. Thus, we sought to understand whether distinct axon classes use unique mechanisms for myelination. To this end, we hypothesized that OLs and their individual myelin sheaths use postsynaptic signaling machinery to coordinate axon identity-dependent myelination.

In this study, we used larval zebrafish to investigate whether Gphn function mediates myelin sheath formation on specific classes of axons defined by neurotransmitter phenotype. We first used transgenic reporters for glutamatergic, GABAergic, and glycinergic neurons to show that each neuronal class is myelinated in the developing spinal cord. Consistent with our hypothesis, Gphn protein localizes to myelin during development and is enriched in sheaths that wrap GABAergic and glycinergic axons. In the absence of Gphn function, myelin sheaths that formed on GABAergic axons were abnormally long, whereas sheath lengths were unchanged on glutamatergic axons. Moreover, with the loss of *gphnb*, overall myelin placement was biased toward glutamatergic axons and away from GABAergic axons. Together, our work illustrates that Gphn plays an important role in selecting inhibitory, GABAergic axons for myelination and point to a paradigm where OLs utilize canonical postsynaptic proteins to establish unique, axon class-specific myelin profiles to coordinate neural circuit function.

## RESULTS

### Glycinergic, GABAergic, and glutamatergic axons are myelinated in the developing spinal cord

As a first step toward investigating axon class-specific myelination, we examined myelination of larval zebrafish spinal cord axons defined by neurotransmitter phenotype. To do this, we used combinations of transgenic reporter genes to simultaneously visualize class-specific axons and myelin. With this approach, we found that glutamatergic axons (Figure 1A), GABAergic axons (Figure 1B), and glycinergic axons (Figure 1C) were myelinated. For each neuronal class, myelinated axons occupied both dorsal (Figure 1 orthogonal views and A’,B’,C’) and ventral spinal cord (Figure 1A’’,A’’’,B’’,B’’’,C’’,C’’’). Additionally, large myelinated glutamatergic and glycinergic axons occupied positions near the midline of the spinal cord (Figure 1A’’,A’’’,C’’,C’’’). Myelinated GABAergic axons were typically smaller in diameter and occupied more lateral positions (Figure 1B’,B’’). Thus, OLs myelinate distinct classes of axons in the dorsal and ventral white matter tracts of the zebrafish larval spinal cord.

**Figure 1.**
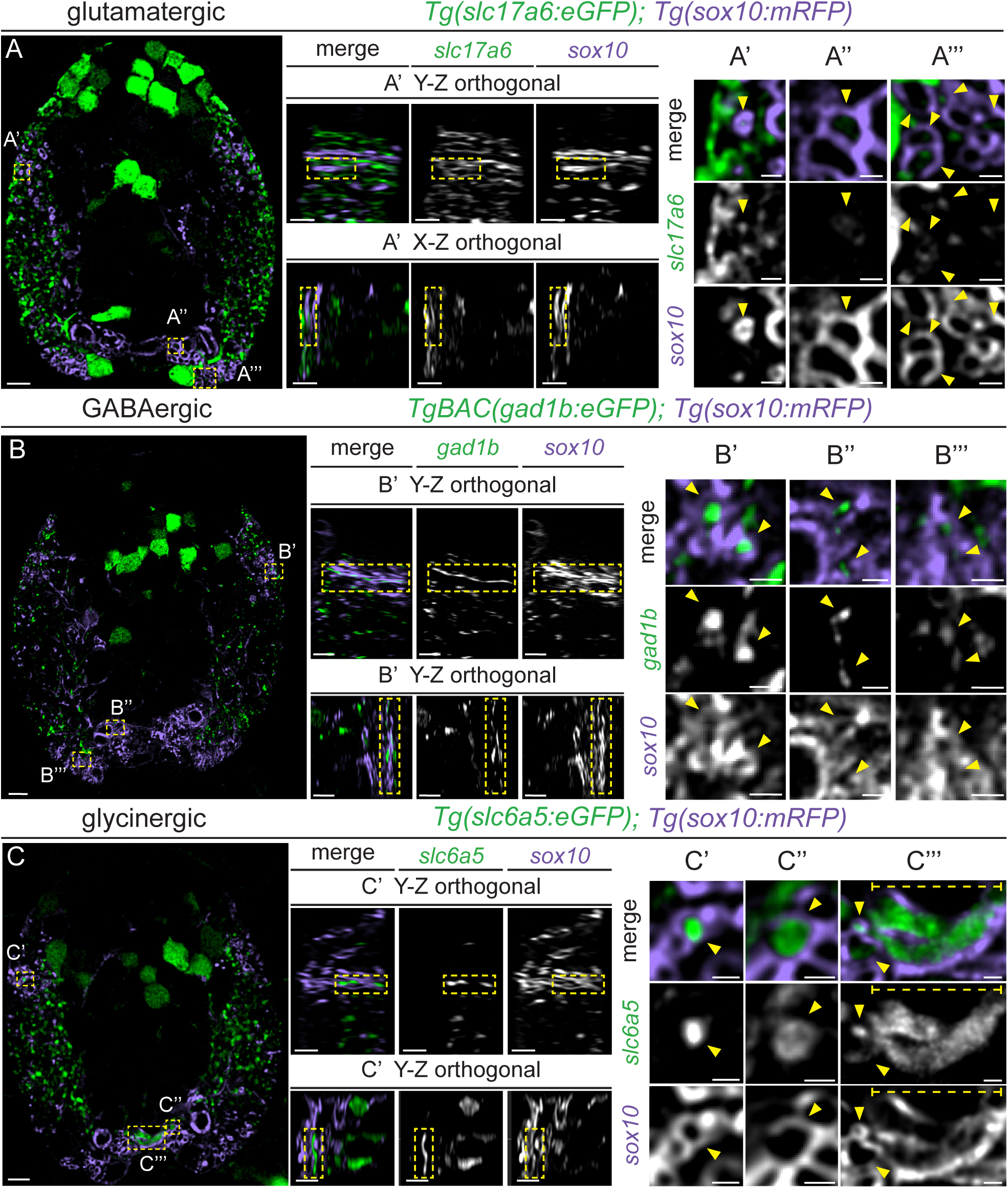
Glycinergic, GABAergic, and glutamatergic axons are myelinated in the developing spinal cord. Representative images of transverse spinal cord sections of 4 dpf larvae expressing the transgene *Tg(sox10:mRFP)* (myelin, purple) along with *Tg(slc17a6:eGFP)* (glutamatergic axons, green, A); *Tg(gad1b:eGFP)* (GABAergic axons, green, B); or *Tg(slc6a5:eGFP)* (glycinergic axons, green, C). (A’-C’’’) Magnification of boxed regions showing individual myelinated axons; (A’-C’) orthogonal views show *sox10*:mRFP^+^ myelin sheaths on a dorsal *slc17a6:*eGFP^+^ glutamatergic axon (A’), a *gad1b:*eGFP^+^ GABAergic axon (B’), and a *slc6a5*:eGFP^+^ glycinergic axon (C’). Magnified transverse views of *sox10:*mRFP^+^ myelin surrounding *slc6a5:*eGFP^+^ glutamatergic axons (A’-A’’), *gad1b:*eGFP^+^ GABAergic axons (B’-B’’), and *slc6a5*:eGFP^+^ glycinergic axons (C’-C’’’) in the ventral spinal cord. Yellow arrows indicate myelinated axons in enlarged panels. A-C scale bars = 5 μm. Zoom scale bars = 2 μm.

### Gephyrin protein localizes to some, but not all, myelin sheaths

OLCs express many genes encoding postsynaptic proteins, some of which appear critical for myelination^24^. Recent work showed that the postsynaptic scaffold protein Gphn localizes to OPC processes^25^ and OLs continue to express *Gphn* at myelinating stages^20–21^. We predicted that if Gphn mediates axon wrapping by myelin membrane then it would occupy nascent myelin sheaths. To test this prediction, we used immunohistochemistry to detect Gphn in transgenic larvae expressing a membrane-tethered myelin reporter (Extended Data 1A-D). This revealed Gphn localization within myelin in both dorsal (Extended Data 1A’,C’) and ventral tracts (Extended Data 1A’’,C’’). Additionally, we determined that the volume of individual Gphn puncta and overall punctal density within myelin increased from 4 dpf to 7 dpf (Extended Data 1E,F). These data show Gphn progressively accumulates in myelin sheaths during development, supporting the notion that Gphn contributes to myelin sheath formation.

To detect Gphn in living animals, we modified a genetically encoded Gphn intrabody^30^ and created a transgene to analyze the sub-cellular localization in OLs at 4 dpf (Figure 2A, Extended Data 2A, fluorescence) and 7 dpf (Figure 2B; Extended Data 2B, fluorescence). Like our findings with Gphn immunohistochemistry, Gphn.FingR puncta per OL increased from 4 dpf to 7 dpf (Figure 2C), an effect that was not influenced by OL sheath number (Figure 2D). At both 4 dpf and 7 dpf, not all sheaths contained Gphn.FingR signal, therefore, to examine sub-cellular localization patterns, we quantified the number of Gphn.FingR puncta per sheath. This showed that individual myelin sheaths had different amounts of Gphn.FingR labeling (Figure 2A’’,B”; Extended Data 2A’’,B’’, fluorescence), with some sheaths containing a high density of puncta (yellow boxes) and others with fewer puncta (magenta boxes). the frequency distribution shifted dramatically between 4 and 7 dpf, reflecting an overall increase in Gphn.FingR expression by 7 dpf (Figure 2E). Notably, some sheaths lacked the Gphn reporter at both 4 dpf and 7 dpf (Figure 2E). These results show that Gphn differentially accumulates in nascent myelin sheaths, raising the possibility that it mediates myelination of distinct classes of axons.

**Figure 2.**
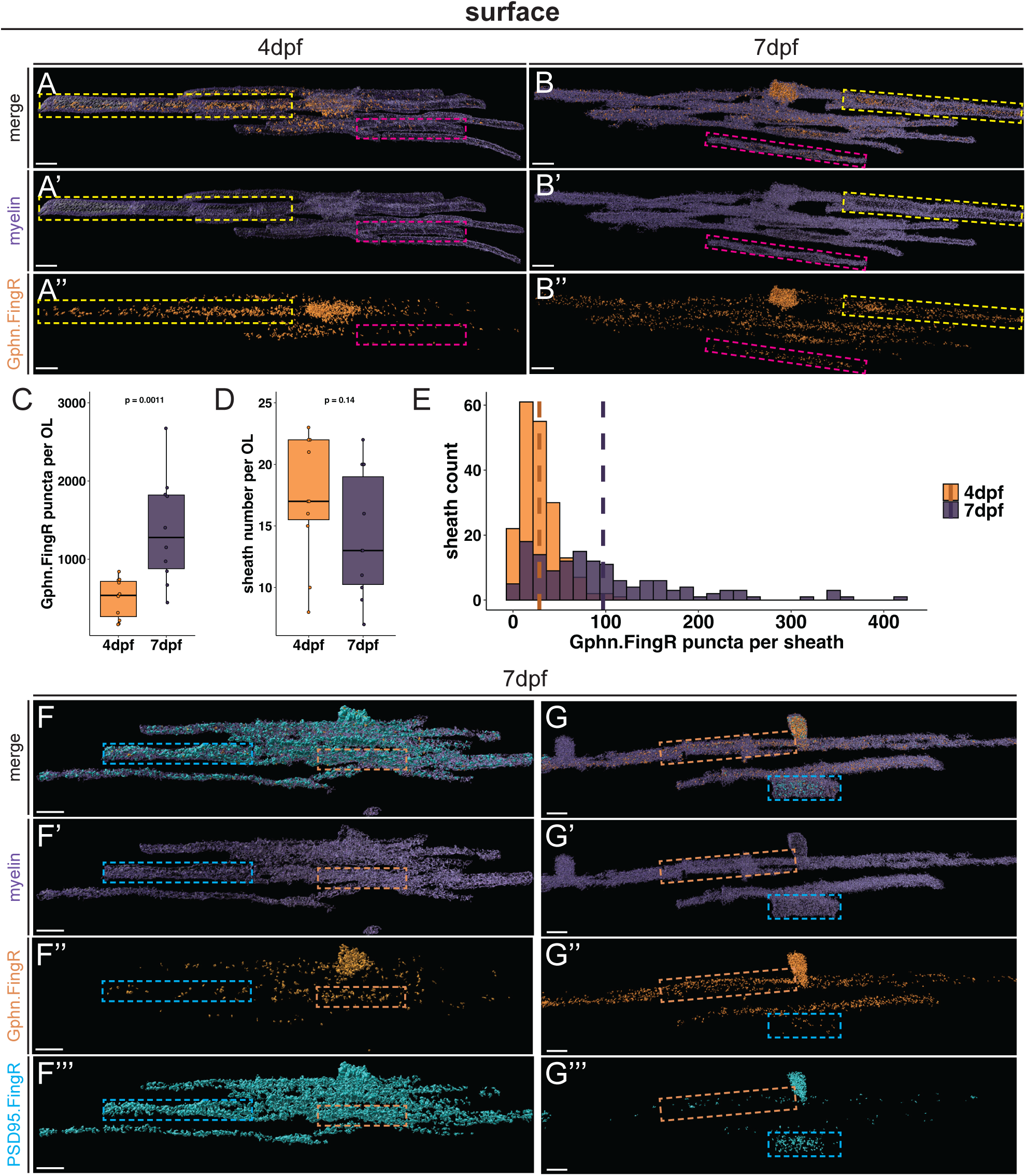
Variable localization of Gphn and PSD95 in myelin sheaths during development. Three dimensional surface models of individual OLs expressing a genetically encoded Gphn intrabody (*zfmbpa:*Gphn.FingR-mScarlet-IL2RGTC-KRAB(A), orange) within *mbpa:*eGFP-CAAX membranes (purple) at 4 dpf (A-A’’) and 7 dpf (B-B’’). Yellow boxes indicate an individual myelin sheath with abundant Gphn intrabody signal, while magenta boxed regions indicate an individual sheath with little Gphn intrabody signal. Quantification of Gphn puncta counts per OL (C; 4 dpf mean = 500±246, 7 dpf mean = 1371±685; Wilcoxon rank sum test) and sheath number per OL (D; 4 dpf mean = 17.5±5.09, 7 dpf mean = 14.1±5.17; Student’s T-Test), at 4 dpf and 7 dpf. p<0.05 is considered significant. (E) Frequency distribution of Gphn puncta per sheath at 4 dpf (orange, n = 11 OLs and larvae) and 7 dpf (purple, n = 10 OLs and larvae), with a dashed line representing the mean (4 dpf mean = 28.5±20.7, 7 dpf mean = 97.2±83.5). (F-G) Surface model of two OLs expressing *mbpa:*tagBFP-CAAX (purple), Gphn intrabody (orange), and the PSD95 intrabody *zfmbpa:*PSD95.FingR-eGFP-CC5TC-KRAB(A) (cyan) at 7 dpf. Cyan boxes highlight sheaths with stronger PSD95 signal, and orange boxes denote sheaths with robust Gphn signal. Scale bars = 5 μm.

Previously, we showed that PSD95, a canonical postsynaptic scaffolding protein at glutamatergic synapses, localizes to nascent myelin sheaths. Because Gphn localizes to GABAergic and glycinergic synapses, the presence of these unique scaffolds in specific OLC processes and myelin sheaths^24–25^ could provide a mechanism for selective, axon class-specific myelination. Therefore, to determine whether Gphn and PSD95 occupy the same or different myelin sheaths we examined OLs that simultaneously expressed Gphn and PSD95 intrabodies (Figure 2F,G; Extended Data 2C,D). At 7 dpf, the density of each type of scaffold varied among myelin sheaths. In particular, some sheaths contained both PSD95 and Gphn puncta, some had more PSD95 puncta than Gphn puncta (Figure 2F’’,G’’; Extended Data 2C’’,D’’, blue boxes), and others had more Gphn puncta than PSD95 puncta (Figure 2F’’,G’’; Extended Data 2C’’,D’’, orange boxes). Altogether, these data indicate that Gphn and PSD95 are not uniformly distributed among newly formed myelin sheaths. Instead, myelin sheaths contain different amounts of these scaffold proteins, supporting the possibility that they are equipped to mediate myelin sheath interactions with distinct axon subtypes.

### Myelin on GABAergic and glycinergic axons has more Gphn than myelin on glutamatergic axons

Because Gphn functions at inhibitory neuronal synapses we predicted that it localizes to myelin sheaths on GABAergic and glycinergic axons. To test this prediction, we investigated Gphn localization within myelin on glutamatergic, GABAergic, and glycinergic neurons marked by transgenic reporter gene expression (Figure 3A,B,C). We used immunohistochemistry to identify Gphn puncta in myelin on axons corresponding to each neuronal class (Figure 3A’,A’’,B’,B’’,C’,C’’, yellow arrows), as well as Gphn puncta in myelin on unmarked axons (blue arrows). This analysis also revealed that some myelin sheaths on specific axonal subtypes lacked Gphn puncta (magenta arrows). Remarkably, myelin on GABAergic and glycinergic axons contained significantly more Gphn than myelin on glutamatergic axons (Figure 3D). This further supports our model that postsynaptic proteins mediate OL interactions with specific classes of axons.

**Figure 3.**
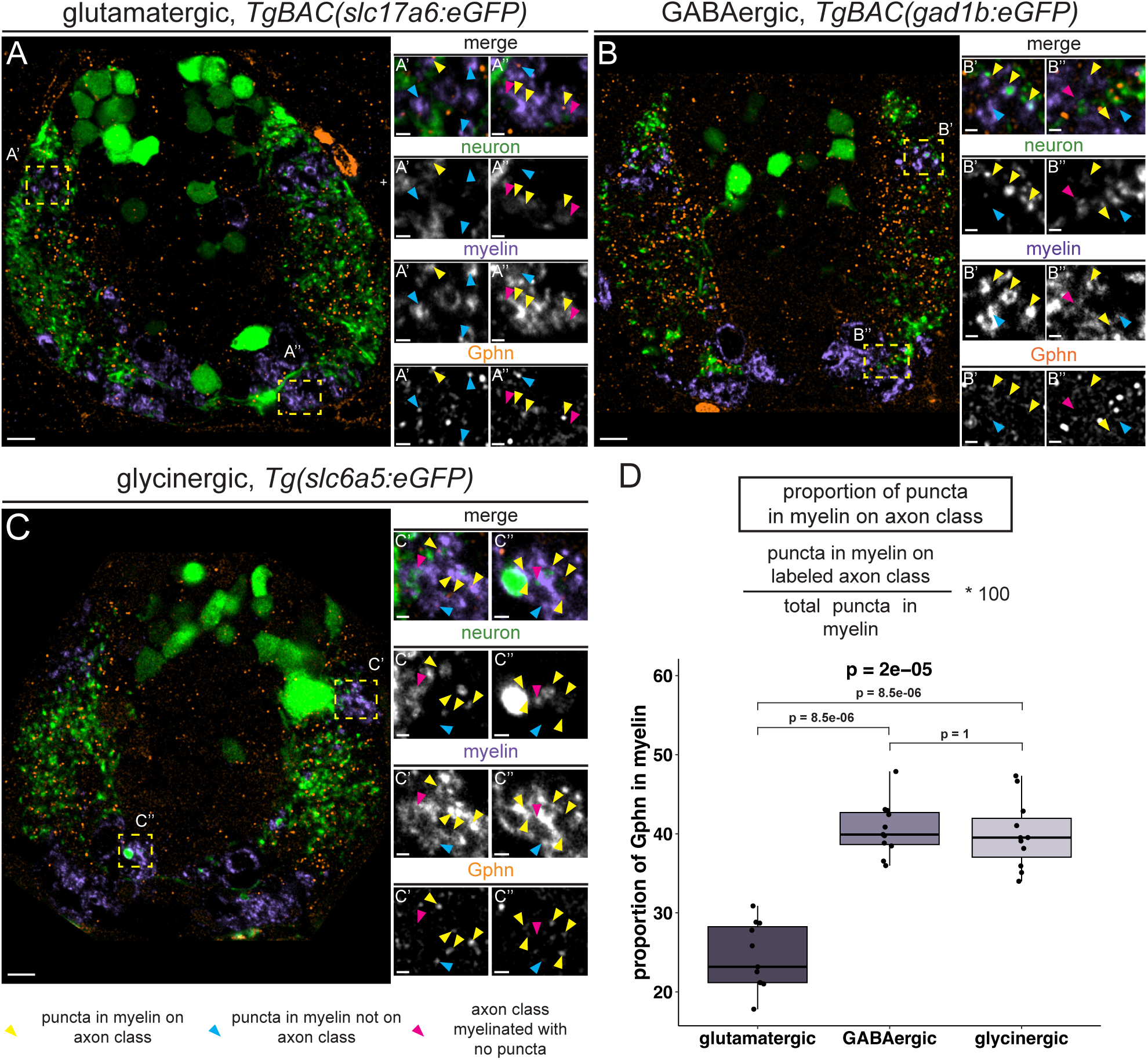
Enrichment of Gphn protein in myelin on GABAergic and glycinergic axons. Representative transverse images of a *Tg(slc17a6:eGFP); Tg(mbp:mCherry-CAAX)* larva (A) *Tg(gad1b:mScarlet-CAAX); Tg(mbp:eGFP-CAAX)* larva (B), and a *Tg(slc6a5:eGFP); Tg(mbp:mCherry-CAAX)* larva (C) processed to detect Gphn at 7 dpf. (A’ and A’’) Magnified panels of boxed regions in A indicating Gphn puncta (orange) in myelin (purple) on glutamatergic axons (green). (B’ and B’’) Magnification of boxed regions in B indicating Gphn puncta in myelin on GABAergic axons (cyan). (C’ and C’’) Magnification of boxed regions in A indicating Gphn puncta in myelin on glycinergic axons (green). In all magnifications, yellow arrows point to Gphn puncta in myelin on the labeled axon class, cyan arrows point to Gphn puncta in myelin not on the labeled axon class, and magenta arrows point to myelin on the labeled axon class without Gphn puncta. (D) Quantification of the percent of Gphn puncta in myelin wrapping each axon class, using the Kruskal-Wallis test for global significance followed by Bonferroni-corrected Wilcoxon multiple comparisons (glutamatergic mean = 24.4%±5.33, GABAergic mean = 40.6%±5.33, glycinergic mean = 39.9%±6.31). p<0.05 is considered significant. All groups n = 11 larvae. Scale bars = 5 μm, zoom panel scale bars = 1 μm.

### Gphn regulates myelin sheath length

To investigate Gphn function in myelination we created loss-of-function *gphn* mutants using CRISPR/Cas9 genome editing. Zebrafish have 2 *gphn* paralogs, *gphna* and *gphnb,* likely from an ancestral genome duplication event^31–32^. By simultaneously targeting *gphna* and *gphnb* we generated 2 lines with mutations in both genes: *gphna^co91^* and *gphnb^co94^, and gphna^co92^* and *gphnb^co95^* (Extended Data 3). Homozygous double mutant larvae do not survive past 4 dpf, likely due to the role Gphn plays in molybdenum cofactor (MoCO) biosynthesis throughout the body^33^. We therefore segregated *gphna* and *gphnb* alleles by outcrossing and then used immunohistochemistry to detect Gphn in larvae with homozygous mutations of each paralog. Whereas *gphna* mutant larvae expressed Gphn in the spinal cord, *gphnb* mutant larvae expressed very little (Extended Data 3D-F). This is consistent with RNA *in situ* hybridization data showing that *gphnb* expression is specific to the central nervous system in zebrafish, whereas *gphna* is globally expressed^34–35^. These data indicate that that *gphnb* mutation mostly eliminates Gphn from the zebrafish nervous system, and we therefore used *gphnb* mutant larvae for our experiments.

To test whether Gephyrin contributes to myelination, we used *mbpa:eGFP-CAAX* expression to label individual OLs in *gphnb* mutant and wild-type larvae and analyzed them at different developmental stages (Figure 4A-B). We focused on dorsal OLs because we could assay all sheath characteristics from individual cells. Neither sheath length nor sheath number differed between homozygous *gphnb* mutant and wild-type larvae at 3 dpf (Figure 4C-E) or 4 dpf (Figure 4C-E). However, by 7 dpf, myelin sheaths were significantly longer in *gphnb* mutant larvae than in wild-type larvae (Figure 4C-E). To confirm that this myelin phenotype is due to loss of Gphnb function and not an off-target mutagenic event, we performed a complementation test using two *gphnb* alleles derived from independent founders (Extended Data 3G). At 7 dpf, there was no difference in sheath length, number, or cumulative sheath length between trans-heterozygous *gphnb^co94/95^* mutant larvae and homozygous *gphnb^co95^* mutant larvae (Extended Data 3H-J). Together, these data indicate that Gphn limits sheath growth.

**Figure 4.**
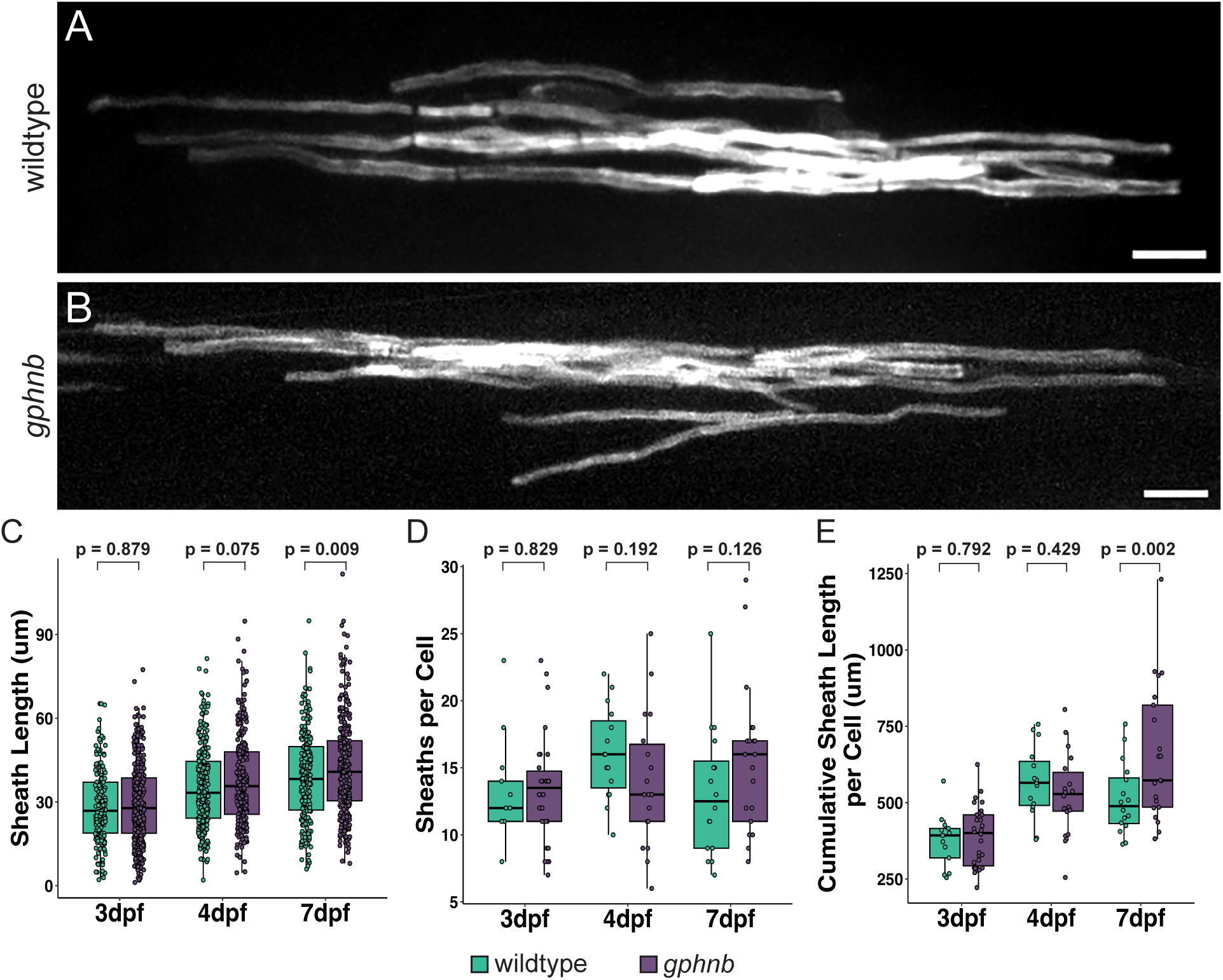
Progressive increase in myelin sheath length in *gphnb* mutant larvae over development. Representative images of mosaically labeled OLs of wild-type (A) and *gphnb* mutant (B) larvae at 7 dpf. Statistical comparisons of sheath characteristics at 3 dpf, 4 dpf, and 7 dpf in wild-type (teal) and *gphnb* mutant (purple) larvae, including individual sheath length (C; 3 dpf, wildtype mean = 28.8 μm±13.6, *gphnb* mean = 29.0 μm±14.0; 4 dpf, wildtype mean = 35.3μm±14.7, *gphnb* mean = 37.7 μm±16.2; 7 dpf, wildtype mean = 39.1μm±16.1, *gphnb* mean = 42.6μm±17.0) total sheaths per cell (D; 3 dpf, wildtype mean = 13.1±3.84, *gphnb* mean = 13.4±3.99; 4 dpf, wildtype mean = 16.1±3.45, *gphnb* mean = 14±4.99; 7 dpf, wildtype mean = 13.1±4.80, *gphnb* mean = 15.4±5.49), and cumulative sheath length per OL (E; 3 dpf, wildtype mean = 376 μm±87.4, *gphnb* mean = 389 μm±98.6; 4 dpf, wildtype mean = 567 μm±117, *gphnb* mean = 528 μm±135; 7dpf, wildtype mean = 513 μm±117, *gphnb* mean = 657 μm±221). 3 dpf: wildtype n = 13 OLs and larvae, *gphnb* n = 30; 4 dpf: wildtype n = 15, *gphnb* n = 18; 7 dpf: wildtype n = 16, *gphnb* n = 21; significance determined by a Type-III sum of squares test followed by Bonferroni-corrected multiple comparisons. p<0.05 is considered significant. Scale bars = 10 μm.

Because neurons express Gphn we performed two complementary experiments to test whether myelin sheath length is limited by Gphn function in OLs. First, we expressed human GPHN in OLs of *gphnb* mutant larvae using *mbpa:GPHN-2A-mApple-CAAX* (Figure 5A,B). This rescued individual sheath length (Figure 5C) and cumulative sheath length (Figure 5E), with no change in sheath number (Figure 5D). Second, we expressed a dominant-negative form of human GPHN, consisting of a single amino acid change (Figure 5F) that disrupts Nlgn2 and GlyR receptor clustering at inhibitory synapses^36^, in OLs of wild-type larvae. Neither sheath number nor cumulative sheath length were different between wild-type control and dominant-negative expressing OLs (Figure 5G-H, J-K). However, OL expression of dominant negative GPHN phenocopied the abnormally long sheaths of *gphnb* mutant larvae (Figure 5I). By comparison, OL expression of wild-type GPHN in wild-type larvae (Extended Data 4A-B) did not change sheath length, number, or cumulative sheath length compared to controls (Extended Data 4C-E). Together, these data provide strong evidence that Gphn functions in OLs to mediate myelin sheath formation.

**Figure 5.**
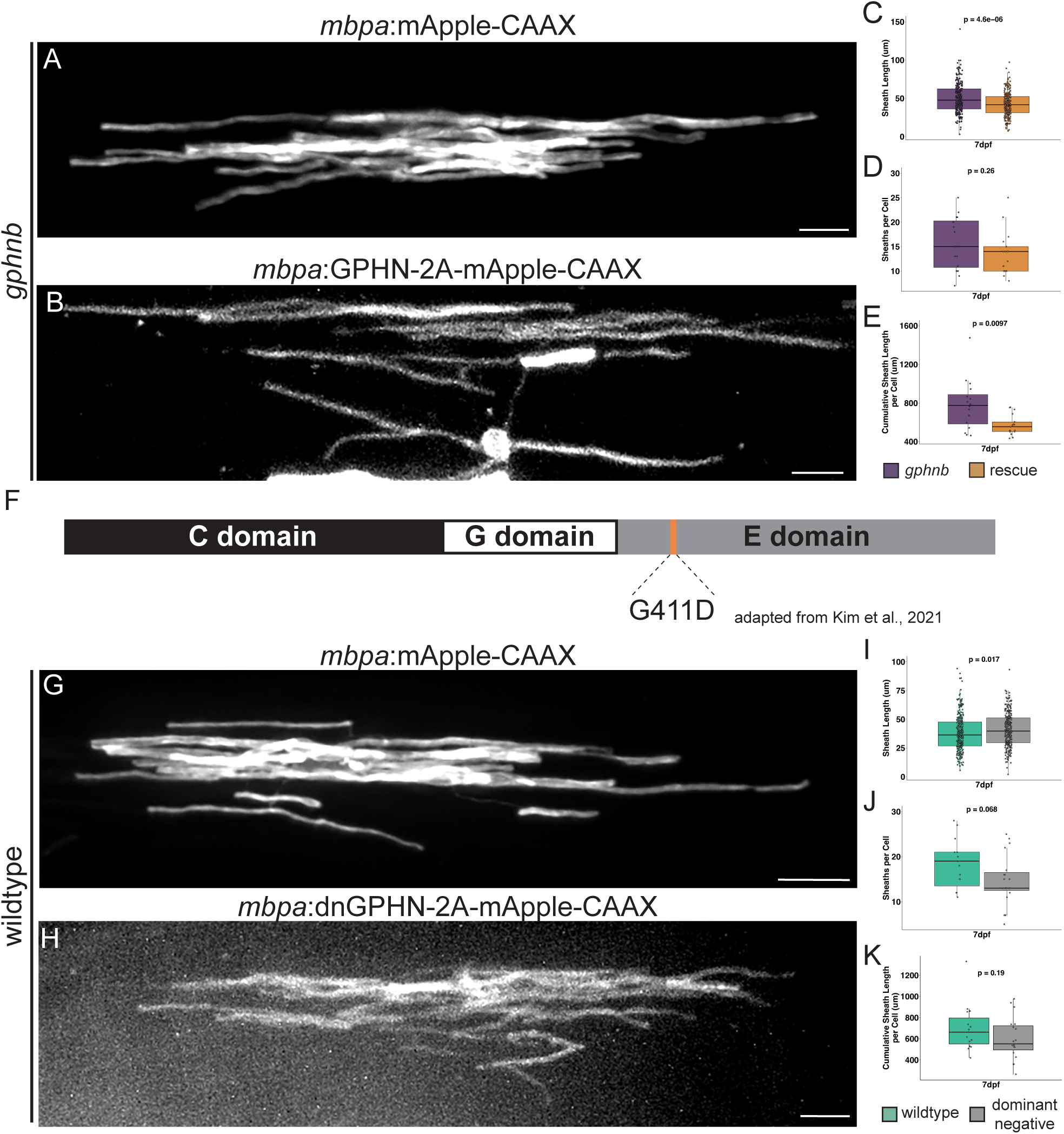
Oligodendrocyte-specific requirements for Gephyrin in myelination. Representative images of individual OLs in *gphnb* mutant larvae expressing control construct *mbpa:*mApple-CAAX (A) and *mbpa:*GPHN-2a-mApple-CAAX (B) at 7 dpf. Quantification of individual sheath length (C; *gphnb* mean = 50.2μm±19.3, rescue mean = 42.1μm±15.7), sheaths per OL (D; *gphnb* mean = 15.6±5.42, rescue mean = 13.6±4.42), and cumulative sheath length per cell (E; *gphnb* mean = 780μm±261, rescue mean = 572μm±104) in 7 dpf larvae. (F) Schematic representing the location (orange bar) and amino acid change in a dominant negative construct adapted from Kim et al., 2021. Example images of OLs in wild-type expressing control plasmid *mbpa*:mApple-CAAX (G) and the dominant negative Gphn construct *mbpa:*dnGPHN-2a-mApple-CAAX (H) at 7 dpf. Quantification of individual sheath length (I; wildtype mean = 38.1μm±15.7, dominant negative mean = 40.7μm±14.6), sheaths per OL (J; wildtype mean = 18.3±5.41, dominant negative mean = 14.7±5.64), and cumulative sheath length per cell (K; wildtype mean = 699μm±223, dominant negative mean = 600μm±198). (A-E) *gphnb* mutant = 16 OLs and larvae, *gphnb* +GPHN = 17; (G-K) wildtype control = 15; wildtype +dnGPHN n = 19. (C, E, and I) statistical comparisons were performed with the Wilcoxon rank sum test; (D, J, and K) statistical comparisons were performed with the Student’s T-Test; p<0.05 is considered significant. Scale bars = 10 μm.

### *gphnb* mutant larvae are hyperactive, but activity does not determine myelin sheath length

Gphn anchors GABA and glycine receptors at inhibitory neuronal synapses. Because inhibition is critical to curtail excitatory output, we predicted that *gphnb* mutant larvae would be hyperactive as a result from reduced inhibition of locomotive circuits. Therefore, we tracked swimming behavior, and at 7 dpf (Figure 6A), *gphnb* mutant larvae covered more distance (Figure 6B), and swam at increased velocity compared to wild-type control larvae (Figure 6C). Because neuronal activity promotes myelination^15,18–19,37–47^ we tested the possibility that neuronal activity drives formation of long myelin sheaths in *gphnb* mutant larvae. To do this, we blocked neuronal activity using tetrodotoxin (TTX), which inhibits voltage-gated sodium channels. We injected TTX or a control solution into the yolk sac and selected paralyzed fish in the TTX group for imaging at 7 dpf (Figure 6D). In wild-type larvae, TTX-induced silencing reduced sheath number and increased individual sheath length, with no change in cumulative length (Figure 6E-F, I-K). By contrast, TTX-induced silencing did not change sheath number, length, or cumulative length in *gphnb* mutant larvae relative to controls (Figure 6G-H, I-K). We conclude that the excessively long myelin sheaths of *gphnb* mutant larvae do not result from elevated neuronal activity. Additionally, these data raise the possibility that loss of Gphn function impairs neuronal activity-dependent modulation of myelin sheath characteristics.

**Figure 6.**
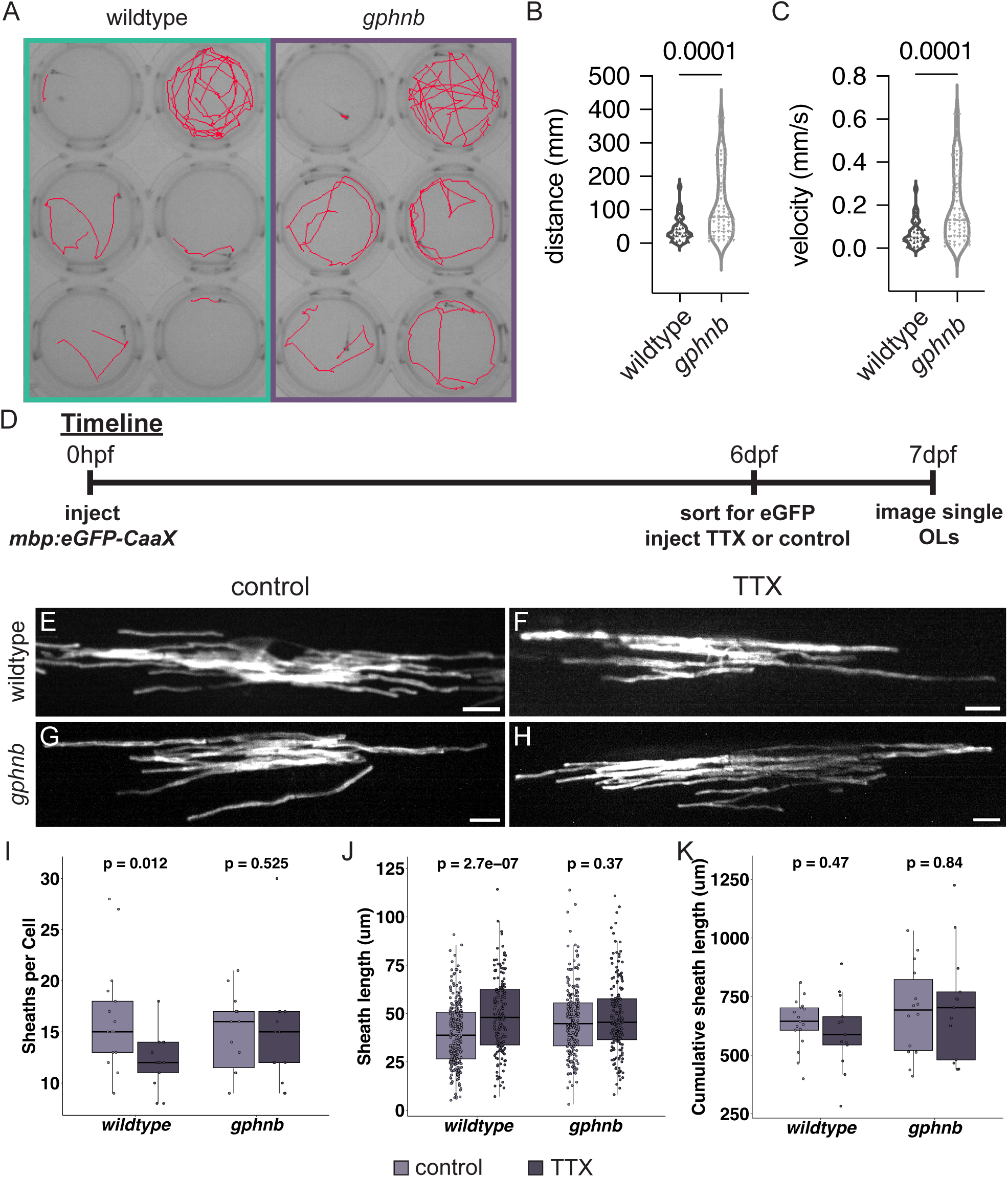
*gphnb* mutant hyperactivity does not account for long myelin sheaths. (A) Tracking data from a behavioral trial with 6 wild-type (left, boxed in teal) and 6 *gphnb* mutant (right, boxed in purple) at 7 dpf. Red lines are traces of the swimming path of an individual larva during the 10 minute video recording. Quantification of cumulative distance swam (B; wildtype mean =, *gphnb* mean =) and swimming velocity (C; wildtype mean =, *gphnb* mean =). (D) Experimental timeline for tetrodotoxin (TTX) experiments: 1) mosaic expression of *mbpa:*eGFP-CAAX at the single-cell stage; 2) eGFP sorting and TTX or control injections at 6 dpf; 3) confocal imaging at 7 dpf. Representative images of OLs in wild-type control (E), wild-type +TTX (F), *gphnb* mutant control (G), and *gphnb* +TTX (H) conditions. Quantification of sheaths per OL (I; wildtype control mean = 16.2±5.14, wildtype TTX mean = 12.2±2.67, *gphnb* control mean = 14.9±3.85, *gphnb* TTX mean = 14.7±5.48), individual sheath length (J; wildtype control mean = 39.3 μm±16.1, wildtype TTX mean = 49.2 μm±19.7, *gphnb* control mean = 46.2 μm±18.1, *gphnb* TTX mean = 48.2 μm±18.9), and cumulative sheath length (K; wildtype control mean = 636 μm ±106, wildtype TTX mean = 598 μm ±159, *gphnb* control mean = 690 μm ±194, *gphnb* TTX mean = 708 μm ±237). B and C comparisons performed with Wilcoxon rank sum tests; wild-type n = 60 larvae; *gphnb* mutant n = 84. I and J comparisons performed with Wilcoxon rank sum tests; K comparisons with unpaired student’s t-tests; wildtype control n = 17; wildtype +TTX n = 13; *gphnb* control n = 14; *gphnb* +TTX n = 13. p<0.05 is considered significant. Scale bars = 10 μm.

### Gphn limits myelin sheath length on GABAergic axons and biases them for myelination over glutamatergic axons

In neurons, Gphn functions specifically at synapses that engage in GABAergic and glycinergic signaling. Could Gphn function similarly in nascent myelin sheaths to mediate specific interactions with GABAergic and glycinergic axons? To test this, we used transgenic reporters to determine whether the sheath length phenotype of *gphnb* mutant larvae is specific to axon class (Figure 7A-D). In wild-type larvae, there was no difference between sheath lengths on glutamatergic and GABAergic axons (Figure 7E). By contrast, myelin sheaths were longer on GABAergic axons than on glutamatergic axons in *gphnb* mutant larvae (Figure 7E). Additionally, there was no difference in the lengths of myelin sheaths formed by individual OLs on GABAergic axons and all other, unlabeled, axons in wild-type larvae (Figure 7F, teal), whereas in *gphnb* mutant larvae the long myelin sheaths of individual OLs were placed on GABAergic axons rather than other, non-labeled axons (Figure 7F, purple). Furthermore, in wild-type larvae the myelin sheaths on glutamatergic axons were slightly longer than those on other axons (Figure 7G, orange), but this difference was absent in *gphnb* mutant larvae (Figure 7G, gray). Together, these data indicate that for individual *gphnb* mutant OLs, long sheaths more frequently occupy GABAergic axons, suggesting an axon class-specific role for Gphn in regulating myelin sheath length.

**Figure 7.**
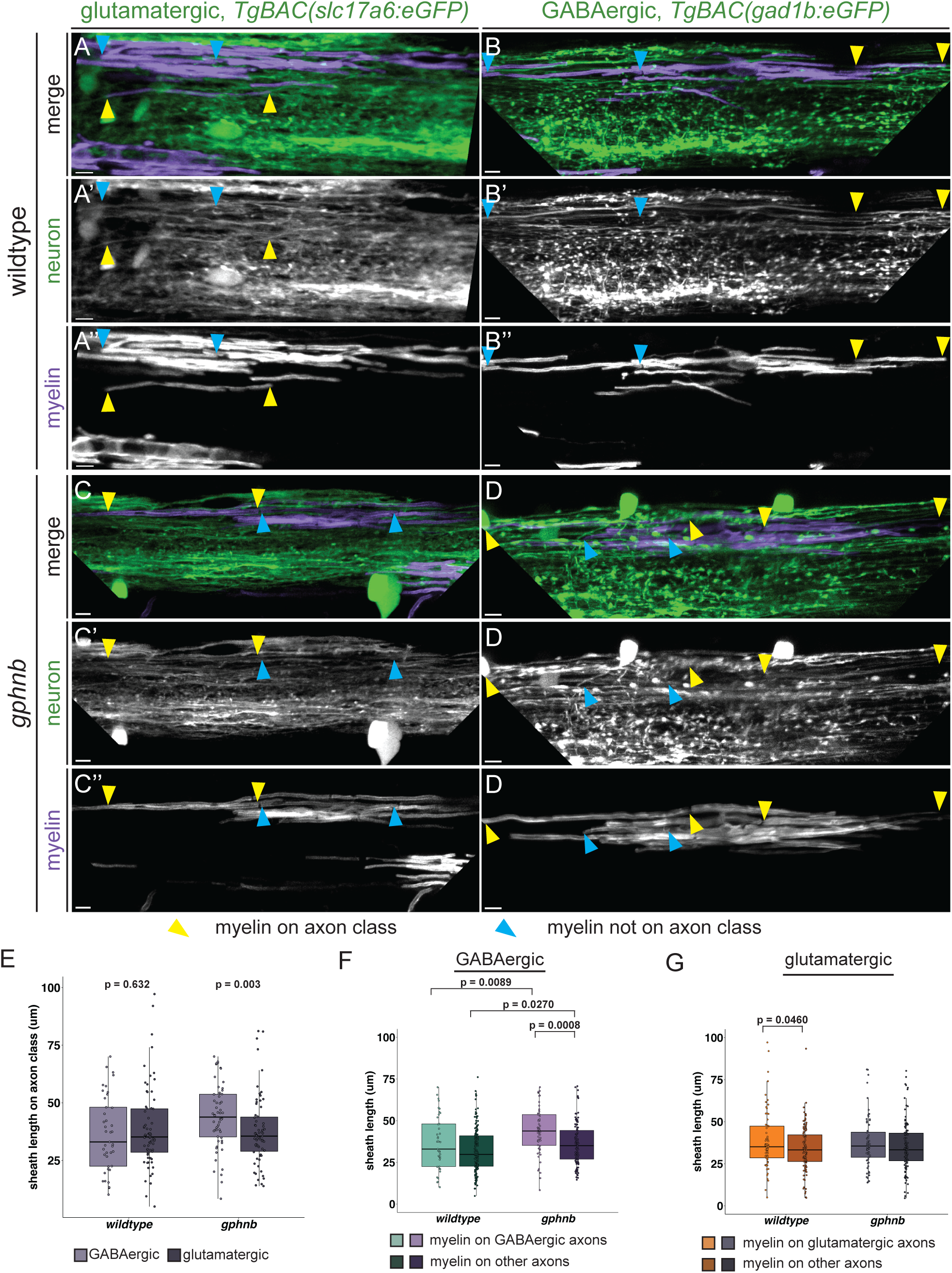
Myelin sheaths are selectively longer on GABAergic axons in *gphnb* mutant larvae. (A-A’’) Representative images of individual OLs mosaically labeled with *mbpa:*mApple-CAAX (purple) in a *Tg(slc17a6:eGFP)* (green) glutamatergic wildtype larva (A-A’’), and *gphnb* mutant (B-B’’); and *Tg(gad1b:eGFP)* GABAergic wildtype larva (C-C’’), and *gphnb* mutant (D’-D”) at 7 dpf. Yellow arrows point to the edges of an individual myelin sheath that wraps the labeled axon class, and cyan arrows point to the edges of an individual myelin sheath that does not wrap the labeled axon class. (E-F) Sheath length quantification of individual sheaths that wrap the labeled axon class between wildtype (teal) and *gphnb* mutants (purple) (E), and comparison of sheath length between axon class (light purple GABAergic, dark purple glutamatergic) within genotypes. (G-H) Comparison of individual sheath length between sheaths that wrap the labeled class and sheaths that don’t wrap the labeled class for GABAergic axons (G) and glutamatergic axons (H). (E) statistical comparisons were performed with the Wilcoxon rank sum test (wildtype GABAergic mean = 36.5 μm±16.3, wildtype glutamatergic mean = 38.7 μm ±17.4, *gphnb* GABAergic mean = 43.9 μm ±14.1, *gphnb* glutamatergic mean = 37.6 μm ±15.2). (F, G) significance determined by a Type-III sum of squares test followed by Bonferroni-corrected multiple comparisons (F, GABAergic: wildtype on axon mean = 36.5 μm ±16.3, wildtype on other axon mean = 32.4 μm ±13.5, *gphnb* on axon mean = 43.9 μm ±14.1, *gphnb* on other axon mean = 36.2 μm ±12.7; G, glutamatergic: wildtype on axon mean = 38.7 μm ±17.4, wildtype on other axon mean = 34.0 μm ±13.5, *gphnb* on axon mean = 37.6 μm ±15.2, *gphnb* on other axon mean = 35.3 μm ±14.6). p<0.05 is considered significant; wildtype glutamatergic n = 10 OLs and larvae; *gphnb* glutamatergic n = 10; wildtype GABAergic n = 12; *gphnb* GABAergic n = 9. Scale bars = 5μm.

Finally, we examined whether Gphn influences which axons are myelinated. Quantification of myelinated glutamatergic (Figure 8A,B) and GABAergic axons (Figure 8C,D) in transverse spinal cord sections showed that *gphnb* mutant larvae had more myelinated glutamatergic axons and fewer myelinated GABAergic axons compared to wild-type control larvae (Figure 8E). These differences were evident in both dorsal (Figure 8F) and ventral spinal cord (Figure 8H), but not among medially myelinated axons (Figure 8G). Notably, there was no difference in the total volume of myelin or myelin volume of dorsal, ventral, or medial myelin between *gphnb* mutant and wild-type larvae (Extended Data 5A-D). These results indicate that the loss of *gphnb* function shifts myelin placement onto glutamatergic axons from GABAergic axons without changing the total amount of myelin. Collectively, we interpret these data to mean that Gphn both biases GABAergic axons for myelination and limits the length of myelin sheaths that form on them, indicating a neurotransmitter identity-dependent function for Gphn in myelination.

**Figure 8.**
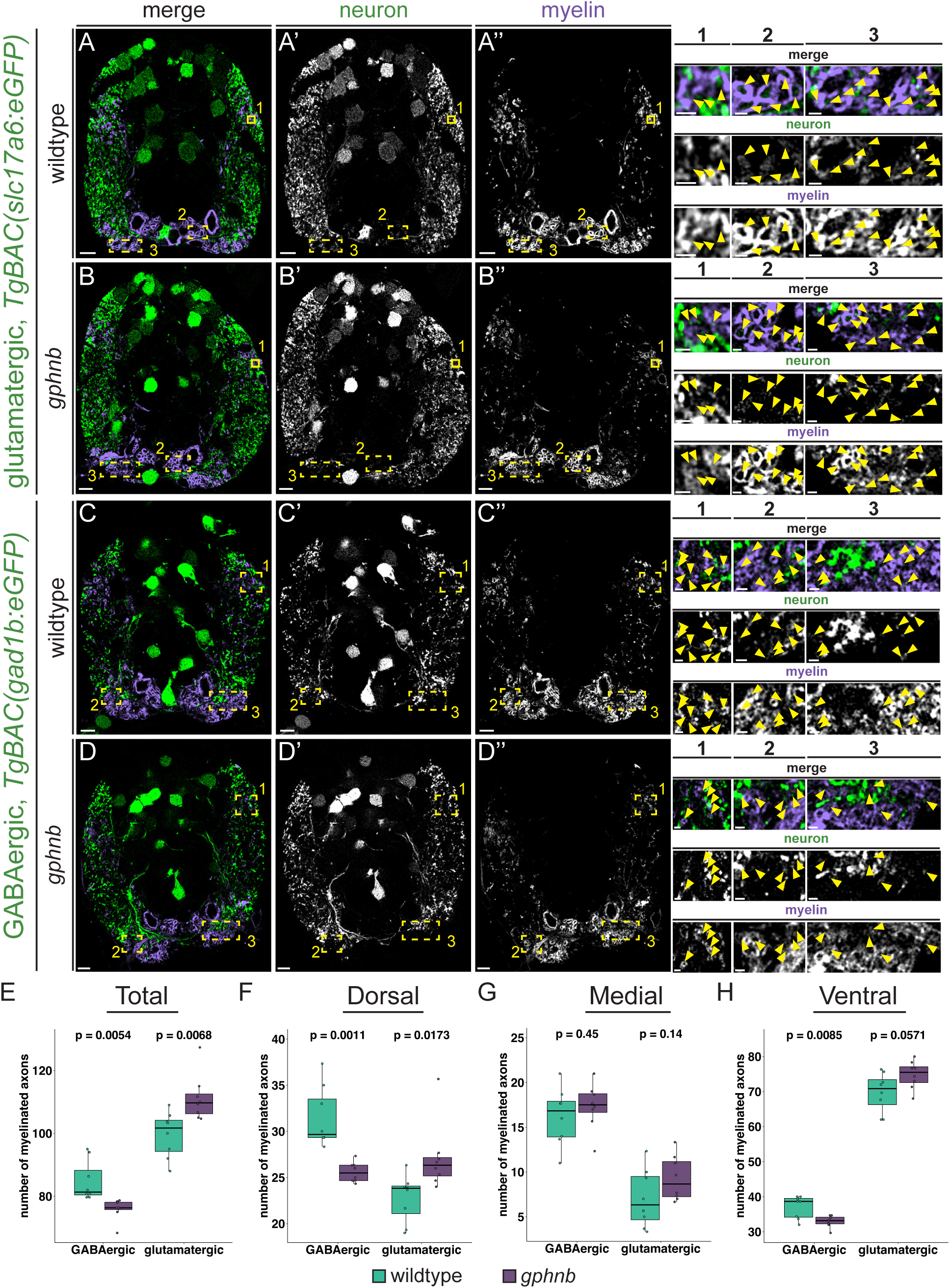
Loss of *gphnb* biases myelin formation onto glutamatergic axons. Representative images of glutamatergic neurons (*TgBAC(slc17a6:eGFP)*, green) and myelin (*Tg(sox10:mRFP)*, purple) transgenic reporters in wildtype (A) and *gphnb* (B) larvae, and GABAergic neurons (*TgBAC(gad1b:eGFP)*, green) and myelin transgenic reporters in wildtype (C) and *gphnb* (D) larvae at 7 dpf. Magnified regions boxed in yellow are enlarged to show dorsal (1) and ventral (2, 3) regions in detail. Yellow arrows point to myelinated axons in magnified panels. Quantification of myelinated axon counts for a single side of the spinal cord for total myelinated axons (E; wildtype GABAergic mean = 84.8±6.39, *gphnb* GABAergic mean = 75.9±3.30, wildtype glutamatergic mean = 99.6±7.26, *gphnb* glutamatergic mean = 111±7.38), dorsal myelinated axons (F; wildtype GABAergic mean = 31.5±3.27, *gphnb* GABAergic mean = 25.6±1.07, wildtype glutamatergic mean = 22.8±2.57, *gphnb* glutamatergic mean = 27.1±3.66), medial myelinated axons (G; wildtype GABAergic mean = 16.2±3.20, *gphnb* GABAergic mean = 17.3±2.56, wildtype glutamatergic mean = 7.04±3.23, *gphnb* glutamatergic mean = 9.29±2.50), and ventral myelinated axons (H; wildtype GABAergic mean = 37.1±3.21, *gphnb* GABAergic mean = 33.0±1.66, wildtype glutamatergic mean = 69.8±5.56, *gphnb* glutamatergic mean = 74.8±3.93). Significance was determined with the Student’s T-Test for E and G, and the Wilcoxon rank sum test for F and H. p<0.05 is considered significant. All groups n = 8 larvae. Scale bars = 5 μm. Magnified panel scale bars = 2 μm.

## DISCUSSION

OLs can myelinate fixed axons and synthetic substrates in vitro^48–50^, indicating that OLs can myelinate axons without the need for specific molecular or cellular cues that distinguish them. However, not all axons are myelinated in vivo and different types of axons are covered by distinct patterns of myelin^10–14^. Furthermore, OLs preferentially place myelin on axons that are more electrically active^15,18–19,43–44,46–47^. These observations suggest that OLs can discriminate between the many different types of axons they encounter in a developing nervous system, but the mechanisms by which they do so remain unknown. In this study, we sought to understand whether OLs use unique molecular machinery to selectively myelinate axons of distinct neurotransmitter classes. Building on prior evidence that synaptic-like mechanisms promote myelin sheath formation, we focused our investigation on Gphn, a scaffolding protein that functions at neuronal postsynaptic terminals that receive inhibitory signals. Altogether, our data support a model where myelinating OLs repurpose classical postsynaptic machinery to facilitate myelination of unique axon classes with specificity characteristic of neuronal synapses (Extended Data 6).

Electrophysiological measurements revealed that neurons make glutamatergic and GABAergic synaptic connections with OL precursor cells (OPCs), which can mediate excitable responses within OPCs^51–56^. Strikingly, calcium activity in OPC processes correlated with Gphn localization, preceding and predicting which processes eventually formed myelin sheaths^25^. Thus, neuronal activity might convey pro-myelinating signals to OPCs via Gphn-mediated synaptic interactions. Whether synaptic mechanisms also mediate axon-OL interactions during myelin sheath formation has been unclear. Although electrical activity was not detected in OLs^57^, activity-dependent calcium transients were evident in nascent myelin sheaths^58–59^. OLs express genes that encode postsynaptic proteins following their differentiation from OPCs^20–21^ and our prior work^24^, together with data we present here, show that the postsynaptic proteins Cadm1b, Caska, PSD95, and Gphn occupy nascent myelin sheaths in zebrafish. Thus, newly differentiating OLs might have the capacity to interact with presynaptic molecules displayed on axons during myelin sheath formation.

Neurons use different molecular complexes to assemble different types of synapses, particularly excitatory and inhibitory synapses. These complexes, anchored by the scaffold proteins Gphn at inhibitory synapses, and PSD95 at excitatory synapses, are necessary for the specificity of neurotransmission. Neurons receive both inhibitory and excitatory inputs by segregating these molecular complexes to different postsynaptic terminals. Consistent with prior observations using mice^12–14^, we determined that OLs myelinate inhibitory GABAergic and glycinergic axons as well as excitatory glutamatergic axons in the zebrafish spinal cord. Remarkably, simultaneous observation of PSD95 and Gphn using intrabodies revealed that they frequently are concentrated in different myelin sheaths of individuals OLs. Moreover, we determined that myelin on GABAergic and glycinergic axons has more Gphn than myelin on glutamatergic axons. These observations raise the intriguing possibility that, similar to neurons, distinct postsynaptic complexes within individual myelin sheaths facilitate interaction with specific types of axons. How then do OLs use postsynaptic factors for myelin sheath formation? Here we learned that in the absence of Gphn function, the myelin sheaths that wrapped GABAergic axons were abnormally long whereas myelin sheaths on glutamatergic axons were unaffected. This effect on sheath length was only evident at 7 dpf, after myelin sheath extension is normally complete in the spinal cord of larval zebrafish^60^. Prior investigations also revealed that disruptions of postsynaptic protein functions have consequences for myelin sheath formation. For example, we showed that OL-specific expression of dominant negative forms of the synaptogenic adhesion proteins Nlgn2b, Cadm1b, and Lrrtm1 caused either longer (Nlgn2b) or shorter (Cadm1b, Lrrtm1) myelin sheaths^24^. Using a transient, cell-type specific CRISPR/Cas9 mutagenesis approach, Li et al. found that knockdown of *dlg4a/b*, which encode PSD95, g*phnb*, and *nlgn3b* in zebrafish reduced the total length of myelin sheaths at 5 dpf^25^. Finally, an OLC-specific γ2-GABA_A_ receptor subunit knock-out resulted in long myelin sheaths in the mouse barrel cortex^26^. This effect was evident on inhibitory, GABAergic parvalbumin (PV)+ interneurons, while neighboring excitatory, glutamatergic spiny stellate cells (SSC) retained normal myelin. Notably, none of these functional manipulations prevented myelination, but they consistently altered myelin sheath length. Neuronal activity also can modulate sheath length^18,43–44^, and sheath length can influence neuronal firing patterns^27–29,61–62^. Thus, postsynaptic proteins expressed by OLs might be key mediators of myelin plasticity, similar to their roles in synaptic plasticity. In support of this possibility, we found that in *gphnb* mutant larvae, myelin sheaths were unresponsive to TTX-mediated inhibition of neuronal activity as they are in wild-type larvae.

Perhaps our most striking finding is that the absence of Gphn function causes myelin sheaths to wrap GABAergic axons less frequently and glutamatergic axons more frequently, without changing the total amount of myelin. With our observation that Gphn is enriched in myelin on GABAergic axons relative to myelin on glutamatergic axons, this suggests that Gphn biases myelination for inhibitory axons, potentially providing a mechanistic explanation for a prior observation that some OLs preferentially myelinate inhibitory axons in the mouse cortex^14^. Together, these results point to a model where postsynaptic proteins function to aid myelin targeting and formation on distinct classes of axons.

An important limitation to our study is that our *gphnb* mutant alleles affect Gphn function throughout the animal. Thus, inhibitory synaptic function likely is impaired in *gphnb* mutant larvae, which could affect myelination. However, our data from several complementary experiments provide strong evidence to support an autonomous role for Gphn in OLs in the process of myelination. A more rigorous approach will require development of a robust and verifiable transgenic method for efficiently eliminating Gphn from OLs. We are currently unable to determine the functional consequences of abnormal myelin caused by OL-specific loss of Gphn. Again, this will require a system for OL-specific mutagenesis.

Collectively, our data reveal a role for canonical postsynaptic proteins in myelination of specific neurotransmitter axon classes. We provide evidence that Gphn biases selection of inhibitory axons for myelin targeting and influences myelin sheath growth. Because myelin can alter axonal conduction velocity^27–29^, myelin targeting and plasticity can provide a powerful influence on circuit development by fine tuning the activity of individual axons within developing neural circuits^63–65^. This is supported by evidence where the specific loss of the γ2-GABA_A_ subunit in OLCs disrupted PV+ firing rate without altering neighboring excitatory SSC frequency^26^. Consequently, the redistribution of myelin patterns on specific axon classes we find in *gphnb* mutant larvae could have important consequences for neural circuit function by disrupting the balance of excitatory and inhibitory signaling (E/I balance)^62–68^. This raises the possibility that genetic variants of synaptic genes that are significantly linked to neuropsychiatric disorders, such as autism and schizophrenia, alter myelination, thereby contributing to the E/I imbalance characteristic of those diseases.

## EXTENDED DATA

**Extended Data 1.**
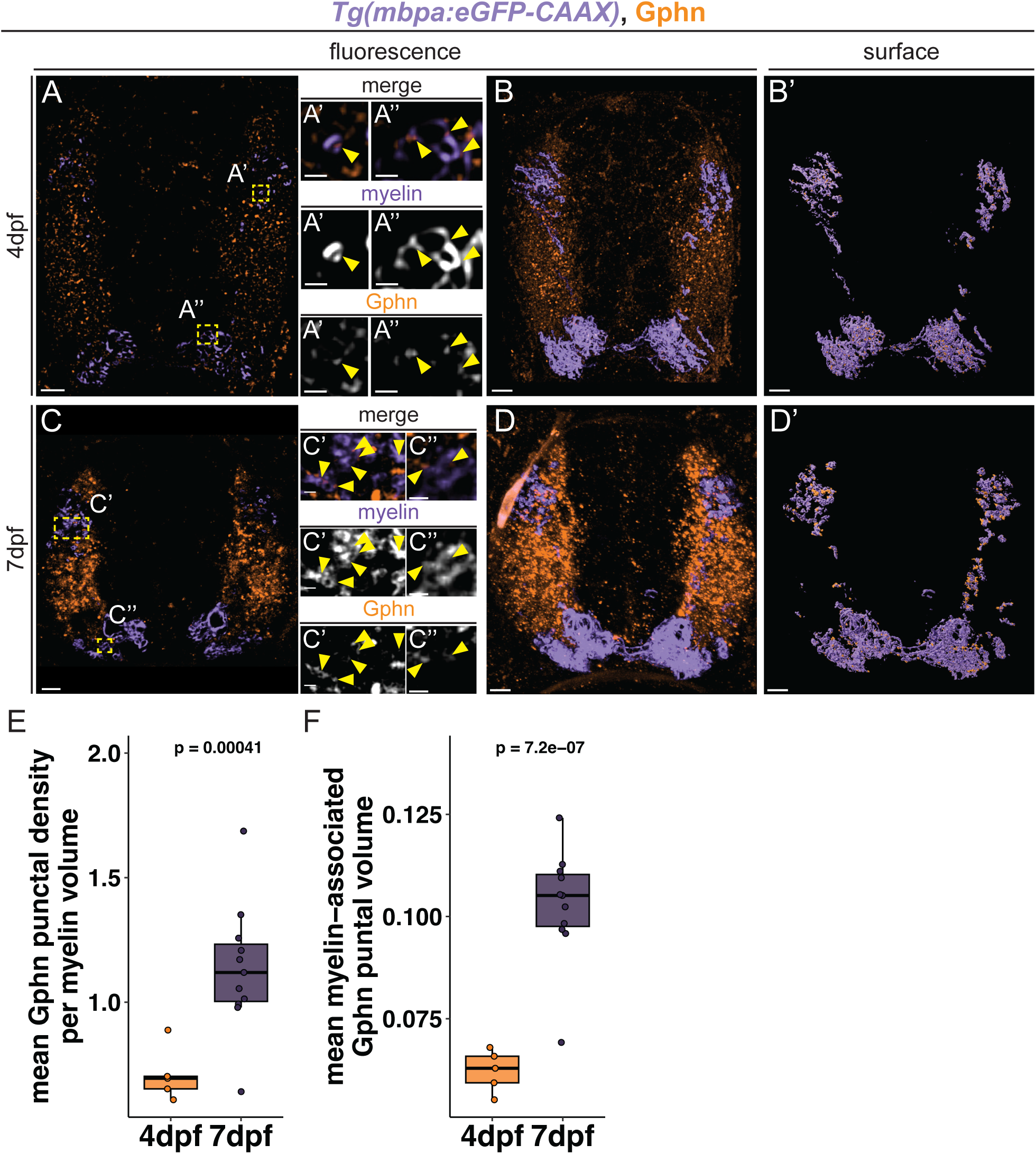
Gephyrin protein localizes to myelin sheaths during development. (A-D) Representative images of the spinal cord in *Tg(mbpa:eGFP-CAAX)* (purple) larvae immunostained for Gphn (green). (A) 4 dpf larva. (A’) Magnification of boxed region in A showing Gphn puncta in dorsal myelin. (A’’) Magnification of boxed region in A of Gphn puncta in ventral myelin. B) 20 μm fluorescent stack of same larva in A. (B’) Surface reconstruction of stack in B of myelin and Gphn puncta contained to myelin channel. (C) 7 dpf larva. (C’) Magnification of boxed regions in C of Gphn puncta in dorsal myelin (C’) and ventral myelin (C’’). (D) 20 μm fluorescent stack of same larva in C. (D’) Surface reconstruction of stack in D of myelin and Gphn puncta filtered by distance to myelin channel ≤ 0 μm. Yellow arrows denote Gphn puncta in myelin. (E) Quantification of the mean Gphn puncta in myelin per 100 μm^3^ at 4 dpf (mean = 0.007±0.002) and 7 dpf (mean = 0.011±0.005). (F) Quantification of the mean Gphn punctal volume (μm^3^) in myelin at 4 dpf (mean = 0.062 μm^3^±0.122) and 7 dpf (mean = 0.102 μm^3^±0.201). Data were compared with unpaired student’s t-tests, where p<0.05 is considered significant; 4dpf n = 5 larvae, 7 dpf n = 11 larvae. A-D scale bars = 5 μm; magnified scale bars = 1 μm.

**Extended Data 2.**
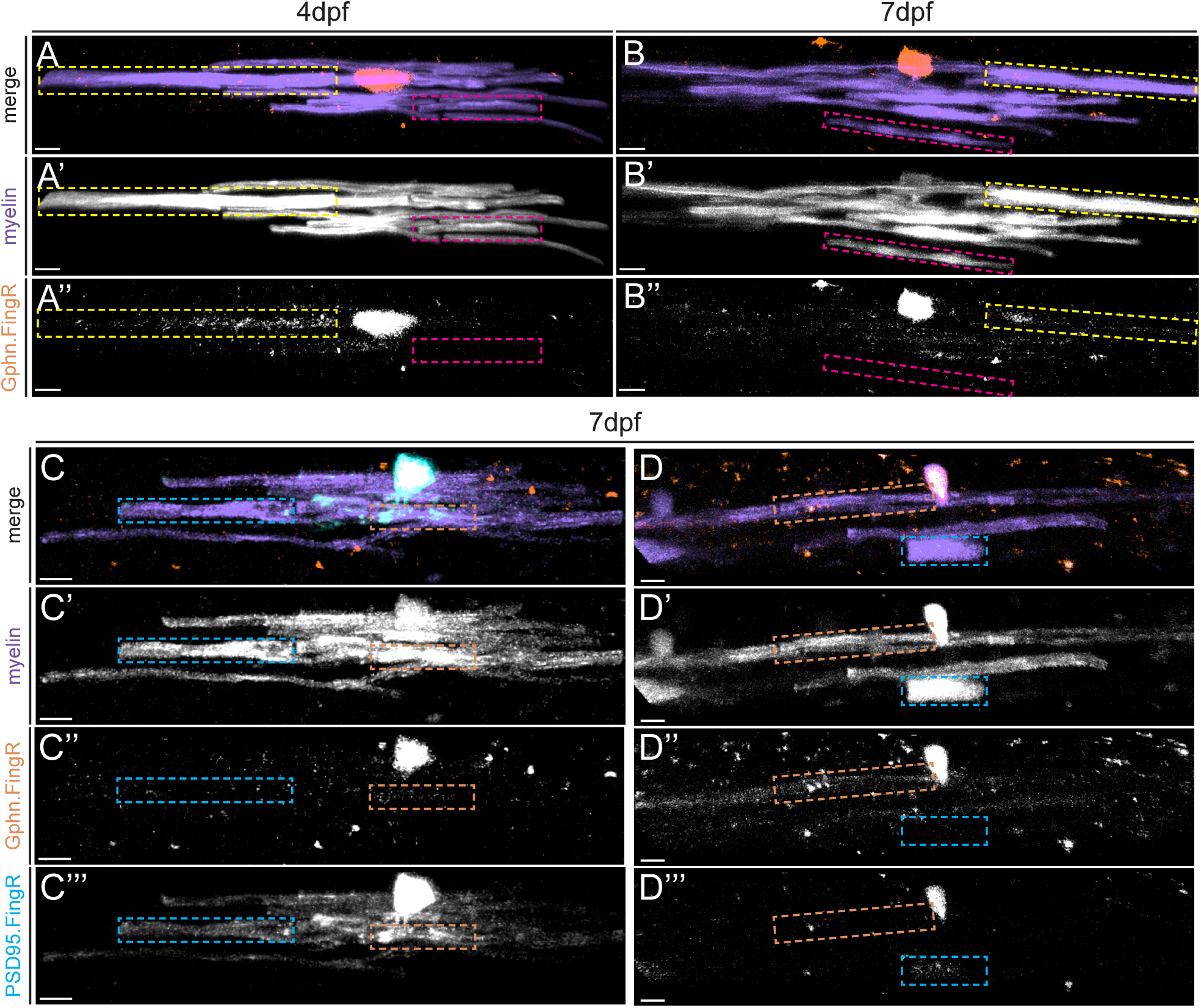
Detection of Gphn and PSD95 in OLs of living larvae using intrabodies. (A) Fluorescent images of the individual OLs expressing *mbpa:*eGFP-CAAX (myelin sheaths, purple, A’) and a genetically encoded Gphn intrabody (orange, A’’) at 4 dpf and 7 dpf (B-B’’). Yellow boxes indicate an individual myelin sheath with abundant Gphn intrabody signal, while magenta boxed regions indicate an individual sheath with little Gphn intrabody signal. (C and D) Fluorescent images of the two OLs in Figure 2E-F, labeled with *mbpa:*tagBFP-CAAX (purple), Gphn intrabody (orange), and PSD95 intrabody (cyan). Cyan boxes indicate individual sheaths with strong PSD95 signal and orange boxes indicate myelin sheaths with robust Gphn signal. Scale bars = 5 μm.

**Extended Data 3.**
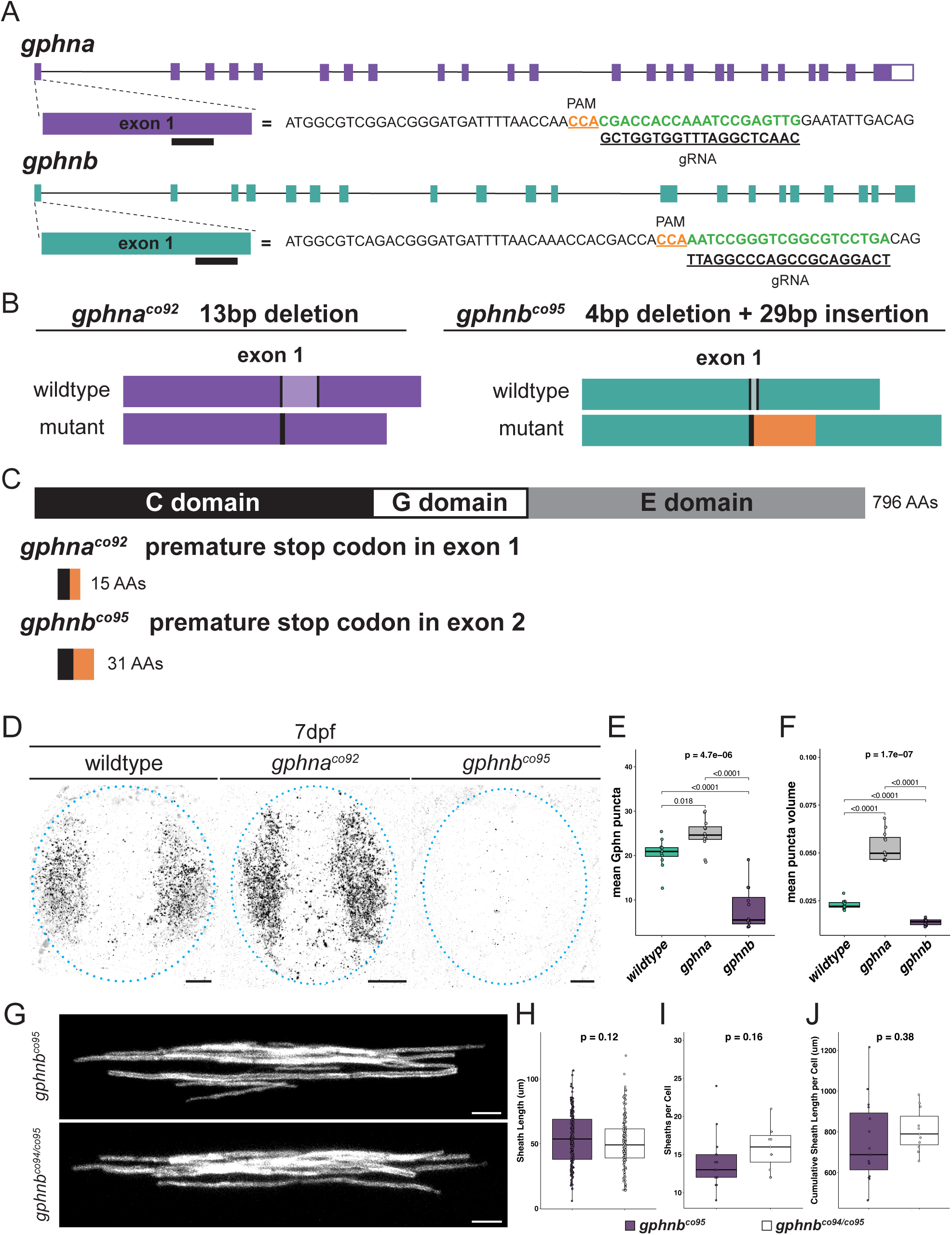
Generation and validation of *gphna^co92^* and *gphnb^co95^* mutant zebrafish. (A) Schematics of genomic sequences for *gphna* and *gphnb*. Exon 1 is enlarged to show where gRNAs align (black bars), and Exon 1 sequence is listed with gRNA sequence (bold and underlined) below the corresponding reverse complement genomic sequence (green) and PAM site (bold and underlined). (B) Sequence changes in exon 1 for *gphna^co92^*(13 bp deletion) and *gphnb^co95^* (4 bp deletion plus 29 bp insertion) alleles. The lighter shaded region between black bars in the wildtype indicates the sequence deleted in the mutants, and the orange region indicates the insertion for the *gphnb^co95^* allele. (C) Gphn protein domain structure with the C domain, G domain, and E domains indicated. Below are schematic representations of the amino acid consequence of the *gphna^co92^* allele (6 missense amino acids followed by a premature stop codon), and the *gphnb^co95^* allele (18 total missense amino acids and a premature stop codon), with altered amino acid regions indicated in orange. (D) Representative images of transverse sections of the spinal cord using immunohistochemistry to detect Gphn in wild-type, *gphna^co92^*, and *gphnb^co95^* larvae at 7 dpf. (E) Quantification of the mean Gphn puncta per 100 μm^3^ volume in D (wildtype mean = 20.5±4.62, *gphnb* mean = 8.00±7.32; *gphna* mean = 24.7±5.39). (F) Quantification of the mean Gphn puncta volume (μm^3^) in D (wildtype mean = 0.023 μm^3^±0.002, *gphnb* mean = 0.014 μm^3^±0.002; *gphna* mean = 0.053 μm^3^±0.008). (E-F) n = 12 larvae each for wildtype, *gphna^co92^* and *gphnb^co95^*; Kruskal-Wallis test was used for non-parametric global comparison followed by Bonferroni-corrected T-Test comparisons). (G) Example images of individual dorsal OLs for *gphnb* complementation test with *gphnb^co95^* and *gphnb^co94/co95^* larvae at 7dpf. Quantification of sheath length (H; *gphnb^co95^* mean = 53.8 μm±19.7, *gphnb^co94/co95^* mean = 51.1 μm±19.1), sheaths per cell (I; *gphnb^co95^* mean = 14±3.66, *gphnb^co94/co95^* mean = 15.8±2.79), and cumulative sheath length (J; *gphnb^co95^* mean = 753 μm±202, *gphnb^co94/co95^*mean = 808 μm±104) for *gphnb^co95^* (n = 15 larvae), and *gphnb^co94/co95^* (n = 11 larvae). Statistical comparisons with Wilcoxon rank sum test (H) and the student’s T-Test (I and J). p<0.05 is considered significant. Scale bars: D = 5 μm; G = 10 μm.

**Extended Data 4.**
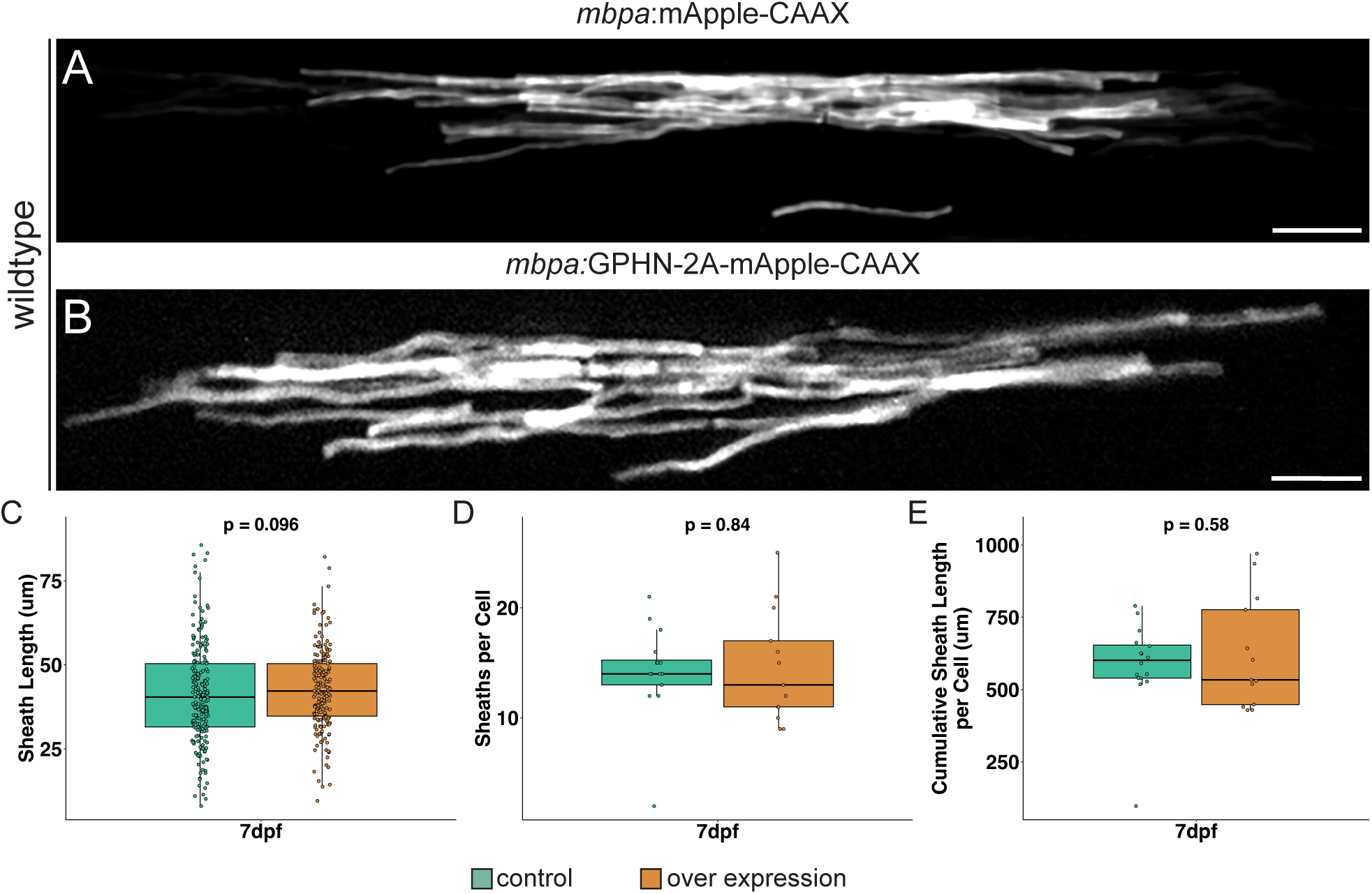
Oligodendrocyte-specific overexpression of human GPHN does not influence myelination in wild-type larvae. Representative images of individual OLs in wild-type larvae expressing control construct *mbpa:*mApple-CAAX (A) and overexpressing GPHN *mbpa:*GPHN-2A-mApple-CAAX (B) at 7 dpf. Statistical comparisons of myelin sheath characteristics in control (teal) and GPHN overexpression (orange), including individual sheath length (C; control mean = 41.2μm±14.7, overexpression mean = 42.7μm±12.0), total sheath count per cell (D; control mean = 14.2±4.09, overexpression mean = 14.5±5.04), and cumulative sheath length per cell (E; control mean = 584μm±154, overexpression mean = 621μm±193). Comparisons were performed with a Wilcoxon rank sum test (C) and unpaired student’s t-tests (D and E), where p<0.05 is considered significant; control n = 13 OLs and larvae; overexpression n = 13. Scale bars = 10 μm.

**Extended Data 5.**
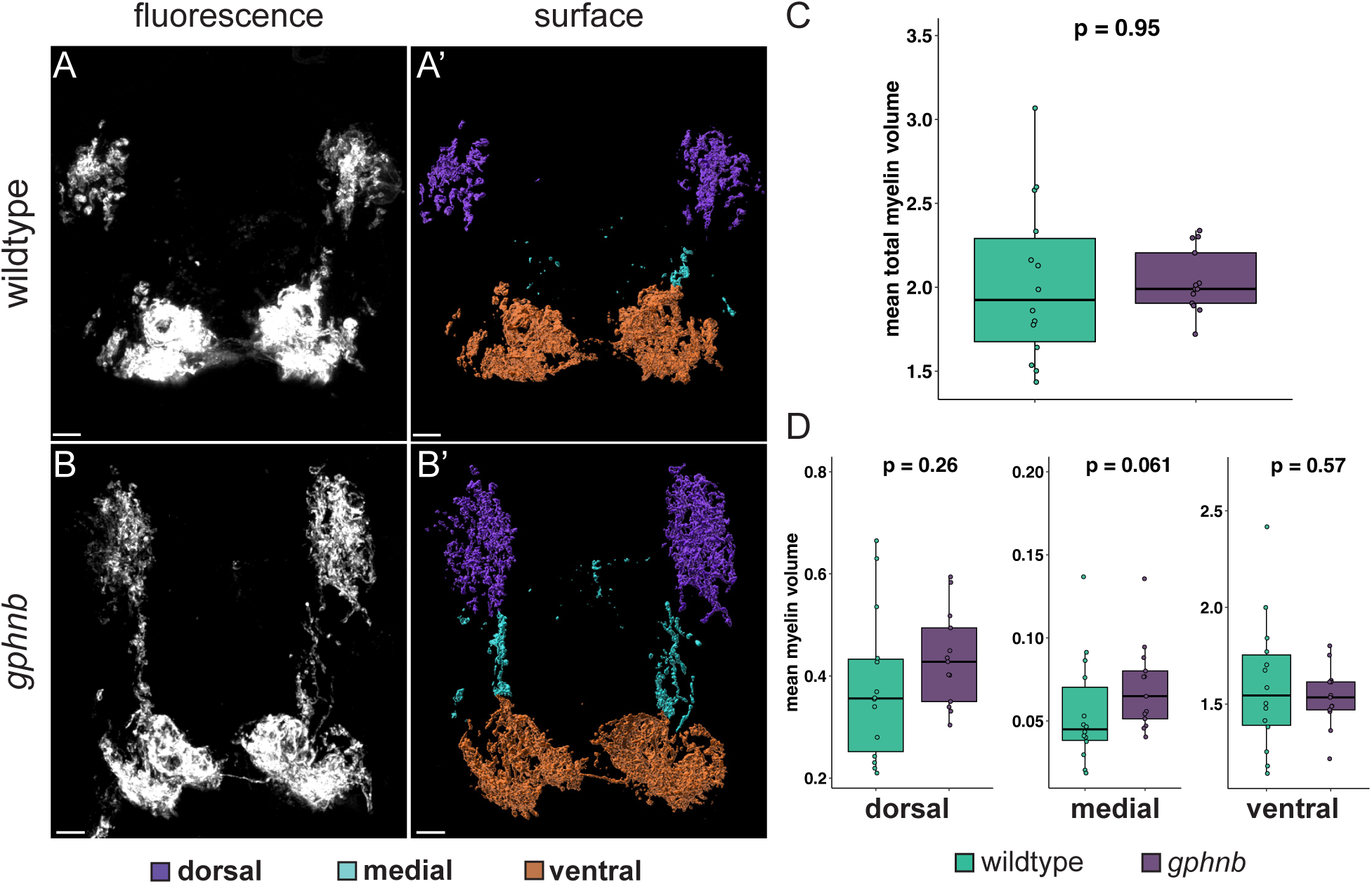
Total myelin volume is unchanged in *gphnb* mutant larvae. Representative images of wild-type (A, A’) and *gphnb* mutant (B, B’) *Tg(mbpa:eGFP-CAAX)* larvae at 7 dpf, including fluorescence projections (A, B) and three-dimensional surface models (A’, B’), with myelin labeled in distinct regions: dorsal (purple), medial (cyan), and ventral (orange). Statistical comparisons of myelin volume at 7 dpf, including mean total myelin volume per 100 μm^3^ in wild-type (teal) and *gphnb* (purple) mutant larvae (C; wildtype mean = 2.23 μm^3^±1.70, *gphnb* mean = 2.04 μm^3^±0.343), and mean myelin volume in dorsal, medial, and ventral spinal cord regions (D; dorsal, wildtype mean = 0.405 μm^3^±0.289, *gphnb* mean = 0.433 μm^3^±0.127; medial, wildtype mean = 0.0567 μm^3^±0.0505, *gphnb* mean = 0.0701 μm^3^±0.0361; ventral, wildtype mean = 1.77 μm^3^±1.41, *gphnb* mean = 1.54 μm^3^±0.262). wildtype n = 14 larvae, *gphnb* = 13 larvae; C and D statistical comparisons were performed with unpaired Student’s T-tests, aside from medial myelin (Wilcoxon rank sum test). p<0.05 is considered significant. Scale bars = 5 μm.

**Extended Data 6.**
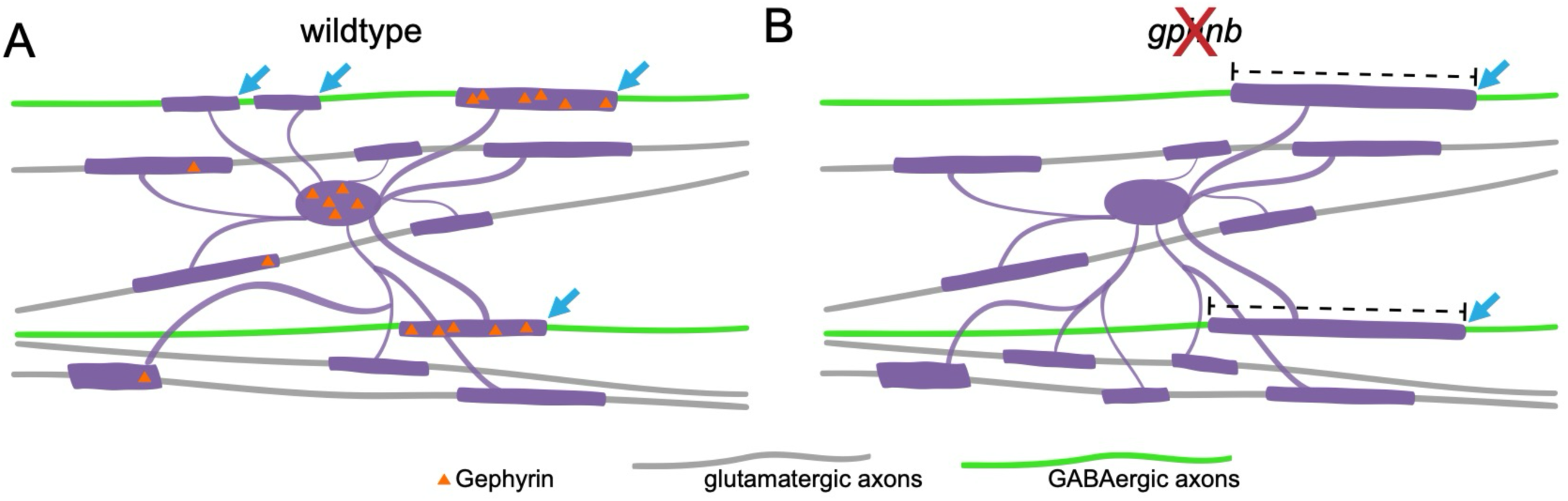
Working model of Gphn function in oligodendrocytes in axon identity-dependent myelination. (A) Gphn localizes to myelin sheaths that wrap GABAergic and glycinergic axons. (B) In the absence of *gphnb*, more glutamatergic axons are myelinated, fewer GABAergic axons are myelinated (blue arrows), and the myelin sheaths that remains on GABAergic axons are longer (dashed line).

## MATERIALS AND METHODS

### Zebrafish Lines and Husbandry

All fish and larvae were handled in accordance with the University of Colorado Institutional Animal Care and Use Committee (IACUC). Previously generated transgenic lines used in this manuscript included: *TgBAC(slc17a6b:eGFP)*^69–70^; *Tg(slc6a5:eGFP)*^71^; *TgBAC(gad1b:eGFP*)^72^; *Tg(sox10:mRFP*)*^vu234,73^*; *Tg(mbpa:eGFP-CAAX)^co58,74–75^*; and *Tg2(mbpa:mCherry-CAAX)^co13,69^*; see key resources table for Zfin IDs). After egg collection or injections, embryos were stored at 28.5°C in petri dishes at a density of roughly 60 embryos per plate or less in egg water (6g instant ocean in 20L miliQ water). Larvae hatched from chorions were stored at similar density in screen media (1X E3, 60μg/mL NaHCO_3_, 145μg/mL CaCl_2_) up to 7 days post fertilization (dpf). For embryo injections, eggs were collected within 30 min of breeding and injected at the single-cell stage. For Tol2 transgenesis injections, 6-25 ng/mL of expression plasmids were injected in 2-3 nL drops. Larvae were then screened prior to imaging for transgenesis reporters or fluorescent protein expression in the spinal cord. When animals for fixed experiments needed genotyping, larvae were anesthetized and quickly tail clipped, and the body was fixed in 4% paraformaldehyde (PFA) while the tail was lysed and used for genotyping. Otherwise, larvae for fixed experiments were euthanized with 4% Tricaine mesylate prior to fixation.

### Mutagenesis

We used the CRISPR/Cas9 system to generate our *gphn* loss-of-function mutations. We used CRISPOR^76^ to design guide RNAs that target the coding sequence of exon 1 for both *gphna* and *gphnb* (figure 5, see key resources table for gRNA sequences). We followed the “Alt-R CRISPR-Cas9 System: *In vitro* cleavage of target DNA with ribonucleoprotein complex” protocol to synthesize each gRNA (http://sfvideo.blob.core.windows.net/sitefinity/docs/default-source/protocol/alt-r-crispr-cas9-protocol-in-vitro-cleavage-of-target-dna-with-rnp-complex.pdf?sfvrsn=88c43107_2, 2017). The RNP complex was created immediately prior to injections with 0.5 μg final concentration of Cas9 (IDT) diluted in Cas9 working buffer (20 mM HEPES, 150 mM KCL, pH7.5), 100 ng each of *gphna* and *gphnb* gRNAs, and incubated at 37°C for 10 min. This solution was then kept at room temperature until injected, with *gphna* and *gphnb* guides co-injected into wild-type embryos with 2-3 nL at the single-cell stage to generate double mutants. Several F0 injected larvae were screened for evidence of insertions and deletions created by DNA cutting and repair with PCR primers that span the guide target site (see key resources table for primers and sequences). F0 larvae were grown to adulthood and individually outcrossed to wild-type fish. Individual F1 larvae were anesthetized and 50 mM NaOH was used for genomic DNA extraction followed by 1M Tris, pH9 neutralization. Mutant allele identification was performed on several individual F1 animals from at least two unique F0 founders each via TA cloning and sequencing followed by alignment with the published genome. F1 adults carrying selected *gphna^co91^*, *gphna^co92^*, *gphnb^co94^*, and *gphnb^co95^*mutant alleles were then outcrossed and F2 generation larvae grown for experiments. All wildtype and mutant larvae used for mutant experiments were genotyped to confirm alleles.

### Behavior

Individual larvae were placed into single wells in a 24-well plate at 7 dpf and given an equal volume of screen media per well. Larvae in 24-well plates were then transferred to the CU Anschutz behavior core in a covered box and allowed to acclimate to the room for 10 min. For the behavior recording, the plate was placed in the DanioVision (Noldus) recording box and larvae were allowed to habituate to the box for 10 min prior to recording start. Larvae were then recorded for 10 min. Following behavior, larvae were anesthetized and lysed for genotyping.

### TTX Injections

For neuronal activity experiments, the voltage-gated sodium channel blocker tetrodotoxin (TTX) was injected into the yolk sac at 6 dpf. All animals were anesthetized with Tricaine and sorted for fluorescent reporter expression stereomicroscope equipped with epifluorescence optics. While anesthetized, larvae were then placed into molds made of 2% agarose in egg water with individual larval wells. Larvae were positioned in the wells such that their yolk sacs faced up, and 3-4 nL buffered TTX (0.6 mM TTX, 0.4 M KCL, 0.05% phenol red, and water) or control solution (0.4 M KCL, 0.05% phenol red, and water) was injected into the yolk sac below the swim bladder. Animals were then removed from the molds and placed into fresh egg water and allowed to recover fully for 24 hrs prior to imaging.

### Cloning

Several plasmids, listed in the key resources table, were generated using the Gateway system and the Tol2 kit^77^. To create the Gephyrin intrabody middle entry vector we used *pCAG_GPHN.FingR-mKate2-IL2RGTC* (Addgene plasmid #46297; RRID: Addgene_46297) and replaced the mKate2 fluorophore with mScarlet *in silico*. Using this plasmid as a template, we custom ordered *pEXPR-zfmbpa:GPHN.FingR-mScarlet-IL2RGTC-KRAB(A)-pA-CC2-Tol2* and a modified version of the PSD95 intrabody we previously published^24^ for co-expression experiments (*pEXPR-zfmbpa:PSD95.FingR-EGFP-CCR5TC-KRAB(A)-pA-CC2-Tol2)* from VectorBuilder (see key resources table for Vector IDs). To create an entry plasmid encoding human GPHN protein, we used plasmid *pECE-M2-GPHN* (Addgene plasmid #31665; RRID: Addgene_31665) and designed primers that introduced a Kozak sequence and attB sites that flank the coding sequence. We amplified human *GPHN* using Phusion polymerase, and then performed a gateway BP reaction with pDONR-221 to create a middle entry *pME-GPHN*. This plasmid was used to create a plasmid to encode dominant negative GPHN as described in Kim et al., 2021^36^. The PASVKDGYAVRAA amino acid sequence at residues 369-381 described in Kim et al., 2021, corresponded to amino acids 405-417 in our *pME-GPHN* sequence. To create the dominant negative mutation in the conserved protein sequence, we generated the amino acid change at G411D *in silico* using SnapGene software (www.snapgene.com) and custom ordered the gene in the pDONR-221 vector from ThermoFisher’s GeneArt custom gene synthesis (figure 5). We also generated middle- and 3’-entry vectors containing *mApple-CAAX* sequence. We used Gateway cloning to create a *pME-mApple-CAAX* vector and InFusion cloning (Takara Bio, 638944) for a *p3E-mApple-CAAX-pA* vector (see key resources table for cloning primers and entry plasmids). The Gateway system was subsequently used to clone these middle-entry vectors and others into expression plasmids with the 5’ entry *p5E-mbpa* plasmid to provide cis-regulatory elements and 3’ reporter fluorophores following a T2A sequence (see expression vectors in key resources table).

### Immunohistochemistry

Larvae were fixed in 4% PFA at each timepoint and washed with 1x PBS after 24 hr. Larvae were then embedded in 1.2% agarose in sucrose, cut to blocks, and soaked in 30% sucrose overnight. Blocks were then dried and frozen on dry ice and stored at −80° until sectioning. Blocks were sectioned on a Leica 1950 cryostat in 20 μm thick sections and thaw-mounted to polarized slides. Slides were mounted in a Sequenza rack followed by immunostaining as previously described^24,74^ (Thermo Scientific, Waltham, MA). Briefly, slides were washed with 1x PBSTx (0.1% Triton-X 100) 3 times for 5 min. Then, a block was added (2% goat serum and 2% bovine serum albumin in 1xPBSTx) for 1 hr. Primary antibodies (see key resources table for antibodies and concentrations) were diluted in block and added to slides and then stored overnight at 4°. The next day, slides were washed in 1x PBSTx every 10 min for 1.5 hr, then secondary antibodies were diluted in block and added and incubated at room temperature for 2 hr (see key resources table for antibodies and concentrations). Secondaries were washed off in 1x PBSTx every 10 min for 1.5 hr, followed by DAPI for 5 min (optional, 1:2000 diluted in 1x PBSTx). For DAPI labeling, slides were washed 3 times for 5 min with 1x PBSTx, followed by mounting. Two drops of Vectashield (Vector Laboratories, H-1000-10) were added followed by application of a No.1 coverslip. Slides were sealed with clear nail polish and allowed to dry at room temperature for 15 min and stored at 4° until imaging. Axon class myelination experiments and total myelin in wildtype and mutant experiments were performed with fixed, transverse sections that were washed with PBSTx and mounted without immunostaining.

### Imaging

#### Live Animal Fluorescent Imaging

Anesthetized larvae sorted for transgenesis reporters or identified as positive for spinal cord fluorescent reporter expression were imaged at 3 dpf, 4 dpf, and 7 dpf. Larvae were mounted in 0.9% agarose with 0.6% tricaine on their sides pressed against a coverslip and imaged on a spinning disk confocal microscope (Carl Zeiss, Oberkochen, Germany) or laser scanning confocal microscope (Carl Zeiss), with a 40x water objective. Individual dorsal OLs were imaged above the yolk sac extension. Standard confocal imaging was used to capture Z-stacks with 0.5 μm step intervals for sheath analysis. Super resolution Airyscan confocal imaging was used on the Zeiss 880 to acquire Z-stacks of individual OLs in *gphnb* and wild-type larvae carrying transgenic neuron reporters to analyze mutant and wild-type sheath length on axon classes with 0.225 μm step intervals. Airyscan imaging was also used to capture all intrabody images with 0.5 μm step intervals. If larvae required genotyping, individual larvae were extracted from the agarose following imaging and lysed with NaOH and Tris and genotyped as described above. Otherwise, larvae were euthanized after imaging.

#### Fixed Tissue Imaging

Transverse imaging was carried out on fixed, 20 μm thick cryosectioned tissue with a Zeiss LSM 880 Airyscan confocal microscope with a 40x oil objective and at least 1.8x zoom. To analyze Gphn localized to myelin on specific axon classes, we used an Andor Dragonfly 200 Spinning Disk Confocal (Oxford Instruments) DMi8 inverted microscope (Leica) with a 63x oil objective. For immunostaining, all images were captured within 2 weeks of staining. For each larva, 3-5 images of the spinal cord were captured serially above the yolk extension and data either summed or averaged by animal. All imaging parameters including Z-step interval, frame size, bit depth, laser power, and gain were kept consistent within experiments.

### Data Analysis

#### Image Blinding

All images were blinded prior to analysis. We used the FIJI (Fiji Is Just ImageJ)^78^, image blinding plugin Lab-utility-plugins/blind-files developed by Nick George and Wendy Macklin (https://github.com/Macklin-Lab/imagej-microscopy-scripts), and only unblinded once analysis was complete.

#### Sheath Analysis

Sheath length was measured using FIJI with the Neuroanatomy plugin SNT (simple neurite tracer)^79^ as described previously^18,24,74,80^. A background subtraction with a rolling ball radius of 50 μm was performed prior to analysis. Briefly, for individual OLs the length of each sheath was traced and length in μm was measured. Each length measurement entry was then counted as an individual sheath and sheaths per cell were summed. Sheath length was plotted for individual sheaths across all OLs analyzed, whereas sheath number and cumulative sheath length were plotted by cell. For sheath length on axon class analysis in wild-type and *gphnb* mutant larvae, images were first Airyscan processed using Zen Black (Carl, Zeiss, version 2.3 SP1). Imaris x64 (Oxford Instruments, 9.9.1-10.2.0) was used to trace sheaths and classify sheaths on reported axons. Sheaths were traced using the Filaments tool, using the manual tracing feature. Individual sheaths were manually traced, and detailed statistics captured for each filament length. Each sheath was then assessed whether it was wrapping reporter axons. Individual sheath length on reporter axons were then plotted by genotype and neuron reporter conditions.

#### Immunohistochemistry analysis

We used Imaris to quantify all immunostaining. With Gphn antibody staining in wild-type and mutant larvae, images were first Airyscan processed for deconvolution in Zen Black. We then performed background subtraction with a 15 μm radius, then cropped the stack to 20 μm. We then created a surface object to quantify puncta with a background subtraction of 0.5 μm, and threshold 5% the maximum value, followed by an intensity-based object splitting and a final volume of ≥0.00125 μm filter. A spinal cord surface was created using the manual creation feature by drawing around the spinal cord on the last slice, duplicating this surface onto the first slice, and creating the surface object from the drawn ROI. The Gphn surface object was then filtered to signal within the spinal cord. We then gathered the specific, detailed volume statistics for total puncta and volume per puncta for each genotype. Three images per larvae were analyzed, with the Gphn puncta measure corrected for spinal cord volume for each image. The mean per fish was then calculated and plotted as puncta per 100 μm^3^.

For Gphn puncta in myelin experiments, images were Airyscan deconvoluted in Zen Black, followed by Imaris background subtraction with a 15 μm radius for the myelin and Gphn channel. Images were then cropped to the inner 20 μm slices from the Z-stack. A surface is created of the myelin channel similar to described above for single OLs, with a background subtraction with diameter of largest sphere of 1 μm, followed by a manually tuned threshold and a split objects filter of 2 μm to best match the fluorescent signal. Extraneous background surfaces were manually deleted. Then, surfaces were created of the Gphn channel as described above in wild-type and mutant conditions. This Gphn surface object was then filtered by distance to the myelin surface with a distance of ≤ 0 μm and duplicated to a new surface. This distance to myelin filter captured the Gphn puncta within the myelin surface. The specific, detailed volume statistics were then generated to provide total puncta and volume per puncta for Gphn in myelin.

To quantify the Gphn puncta in myelin on axon classes, deconvolution was performed in Imaris using 10 iterations, followed by a background subtraction with a 15 μm radius for Gphn channel. We then created myelin and Gphn surface objects as described above for the Gphn in myelin analysis. Once puncta in myelin were duplicated to a new surface object, the slice view option was used with the neuron and myelin fluorescent channels toggled on, and each punctum was manually assessed for whether it is in myelin on the reporter axon. Puncta determined to be in myelin on the reporter axon were selected and duplicated to a new surface object and the specific, detailed volume statistics were generated. The proportion of puncta in myelin on axon class was then calculated by totaling the puncta in myelin on the reporter axons and dividing by the total puncta in myelin (figure 3D).

#### Intrabody Analysis

Intrabody images were Airyscan deconvolution processed and analyzed in Imaris. Background subtraction with a 15 μm radius was performed, then images were cropped to isolate single, dorsal OLs. A surface was created of the OL with a background subtraction with diameter of largest sphere of 1 μm, followed by a threshold that was manually tuned to best match the fluorescent signal. The split objects filter of 2 μm was also used to capture the OL surface more precisely. Intrabody surfaces were created using a background subtraction with a diameter of largest sphere of 0.35 mm, and the threshold was manually adjusted to best match fluorescent signal. The split objects filter of 0.5 mm was used, followed by a voxel filter to further eliminate creation of any background surfaces. The intrabody signal was then filtered to within the OL surface, and any puncta where majority of the volume resided outside of the OL surface with minimal volume inside the surface, and puncta just touching the outside of the OL surface were manually deleted. Puncta in individual sheaths were then manually duplicated to new surface objects and the detailed, specific statistics were used for puncta per sheath quantification. Total puncta per OL was calculated with the summation of the puncta in individual sheaths. This eliminated the bright, self-regulating expression in the cell body.

#### Myelinated axon counts analysis

Transverse images were processed and analyzed in Fiji. First, a background subtraction with a 15 μm rolling ball radius was performed. Then, we created a new sub-stack of the inner 20 μm slices and used the multi-point tool to identify every myelinated axon on one side of the spinal cord. These counts were totaled and separated by region (dorsal white matter tract, medial white matter, and ventral white matter tract). We analyzed 3 images per larvae that were then averaged. Single 0.5 mm sections were used for representative images and the same region was isolated for magnified views between wild-type and *gphnb* mutant larvae.

#### Behavior analysis

The DanioVision software (Noldus) recorded 10-min videos and used the EthoVision TX to automatically track the movement of individual larvae. Total distance moved and velocity were generated and used for analysis between wild-type and *gphnb* mutant larvae.

#### Statistical Analysis

All data analysis, statistics, and plotting were performed using R in R Studio (v 2024.04.2+764), except for behavior analysis which was performed in GaphPad Prism (version 10). Data and statistical analysis were performed using the ggpubr and emmeans packages. Plots were generated with the dplyr and ggplot2 packages. Data from each experiment were tested for normality with the Shapiro-Wilkes test. If data were normally distributed (p>0.05), parametric tests were used to compare groups. If data were not normally distributed, non-parametric tests were used. For tests with two groups, the student’s T-Test or Wilcoxon rank sum test were used. For datasets with multiple comparisons, global significance was tested with a one-way Anova, Kruskal-Wallis test, or a type III two-way ANOVA for unbalanced design, followed by Bonferroni corrected pair-wise T-Tests or Wilcoxon rank sum tests. All experiments were performed by 7 dpf when sex is not yet determined in the zebrafish.

## Key resources table

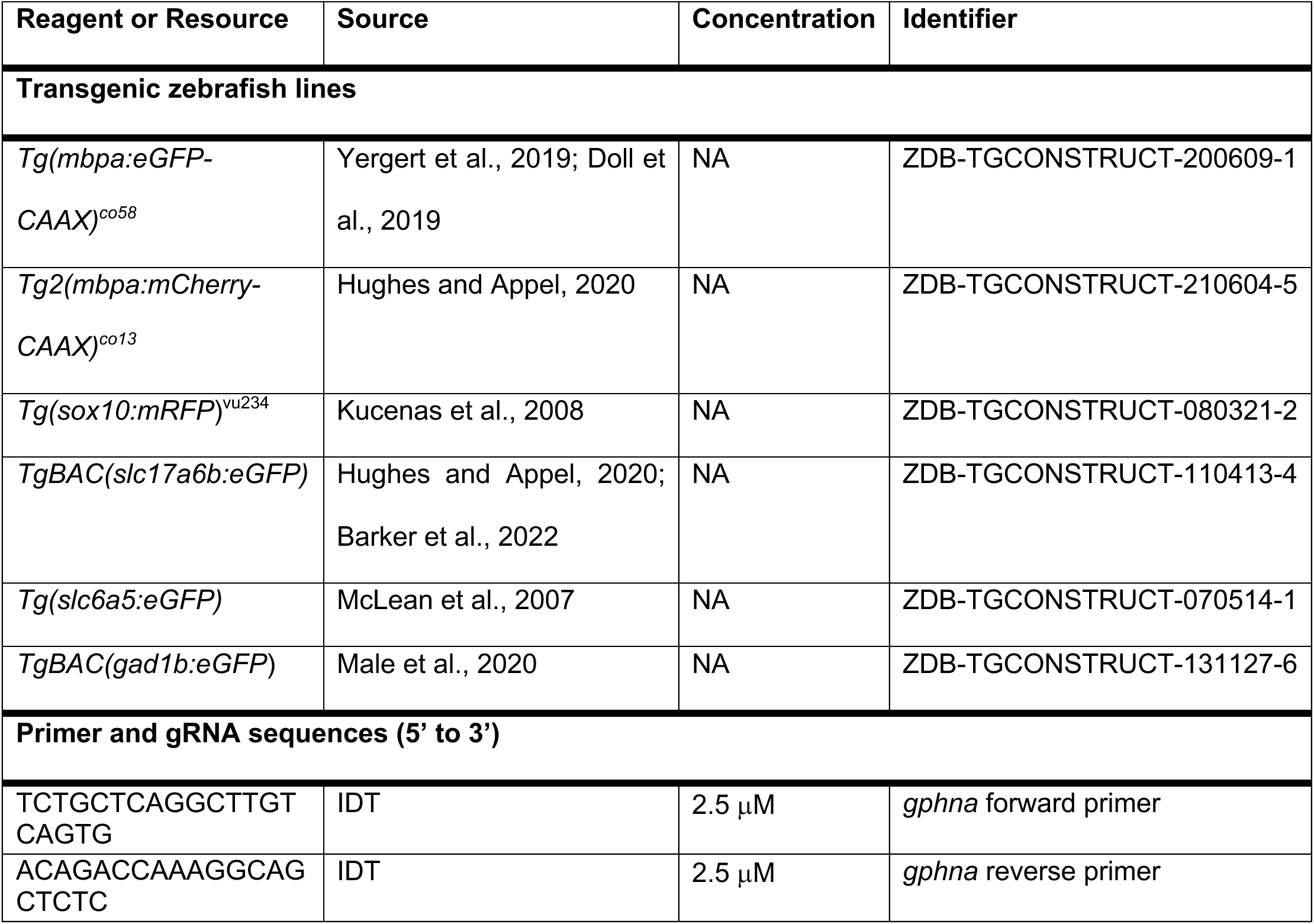
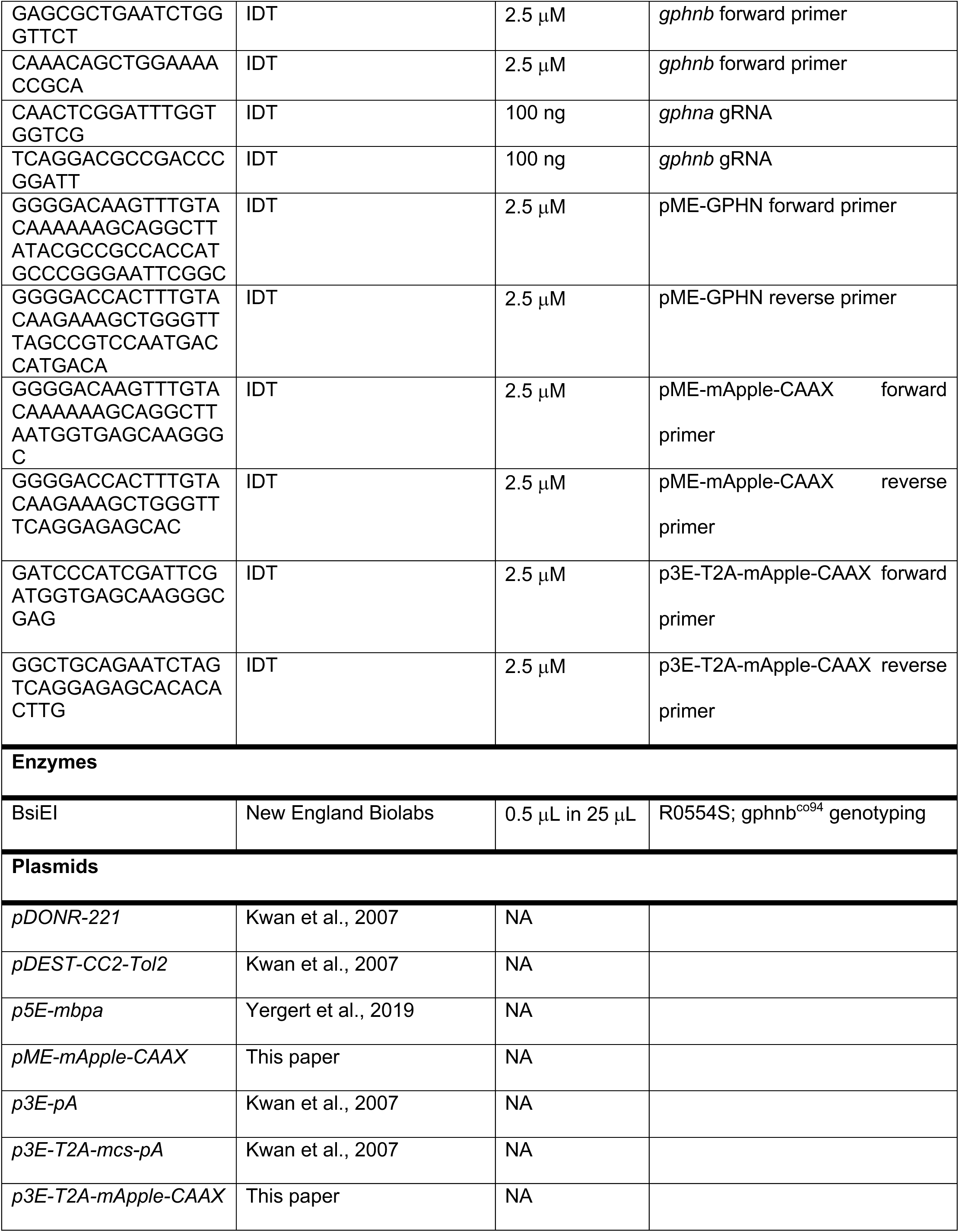
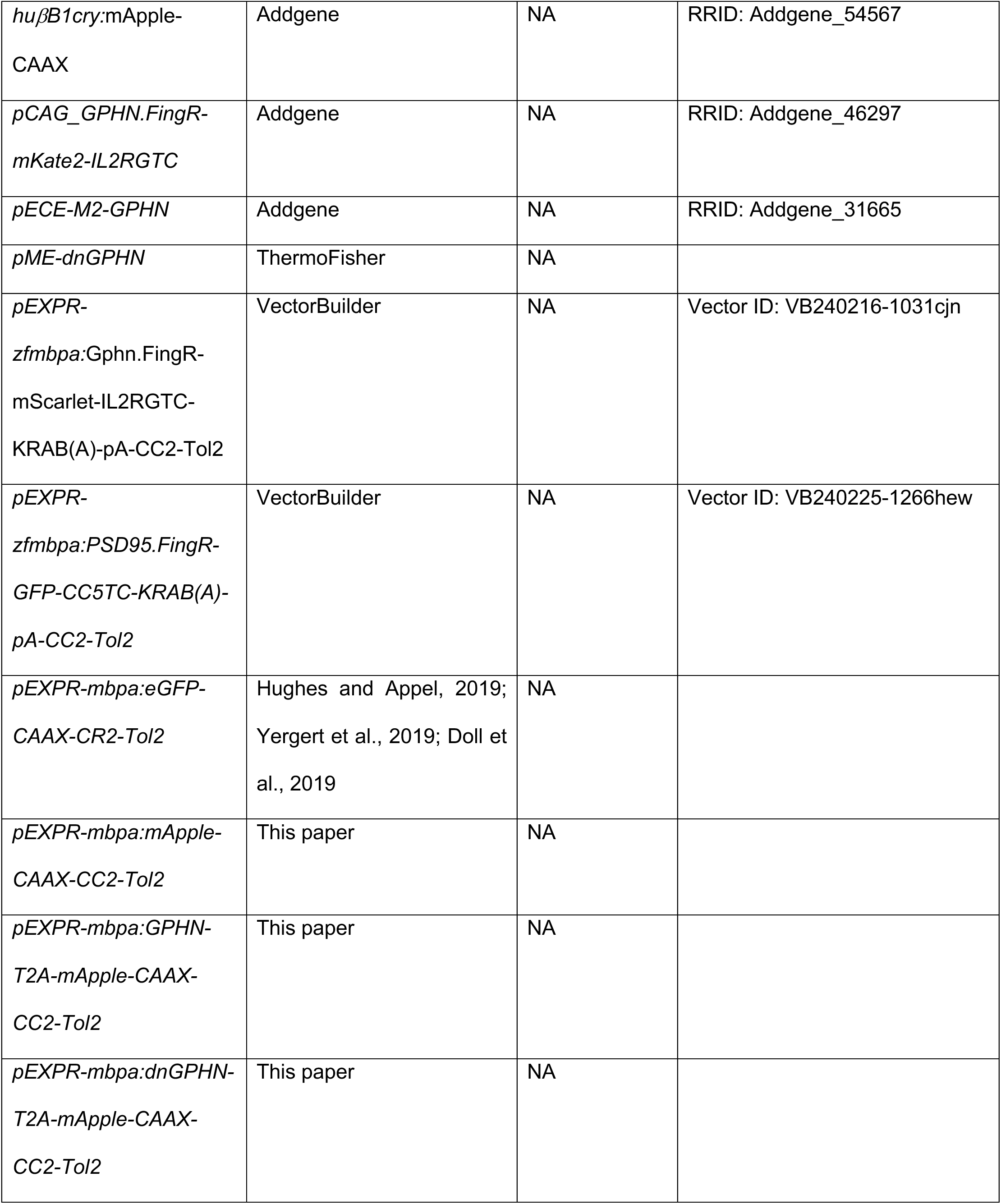
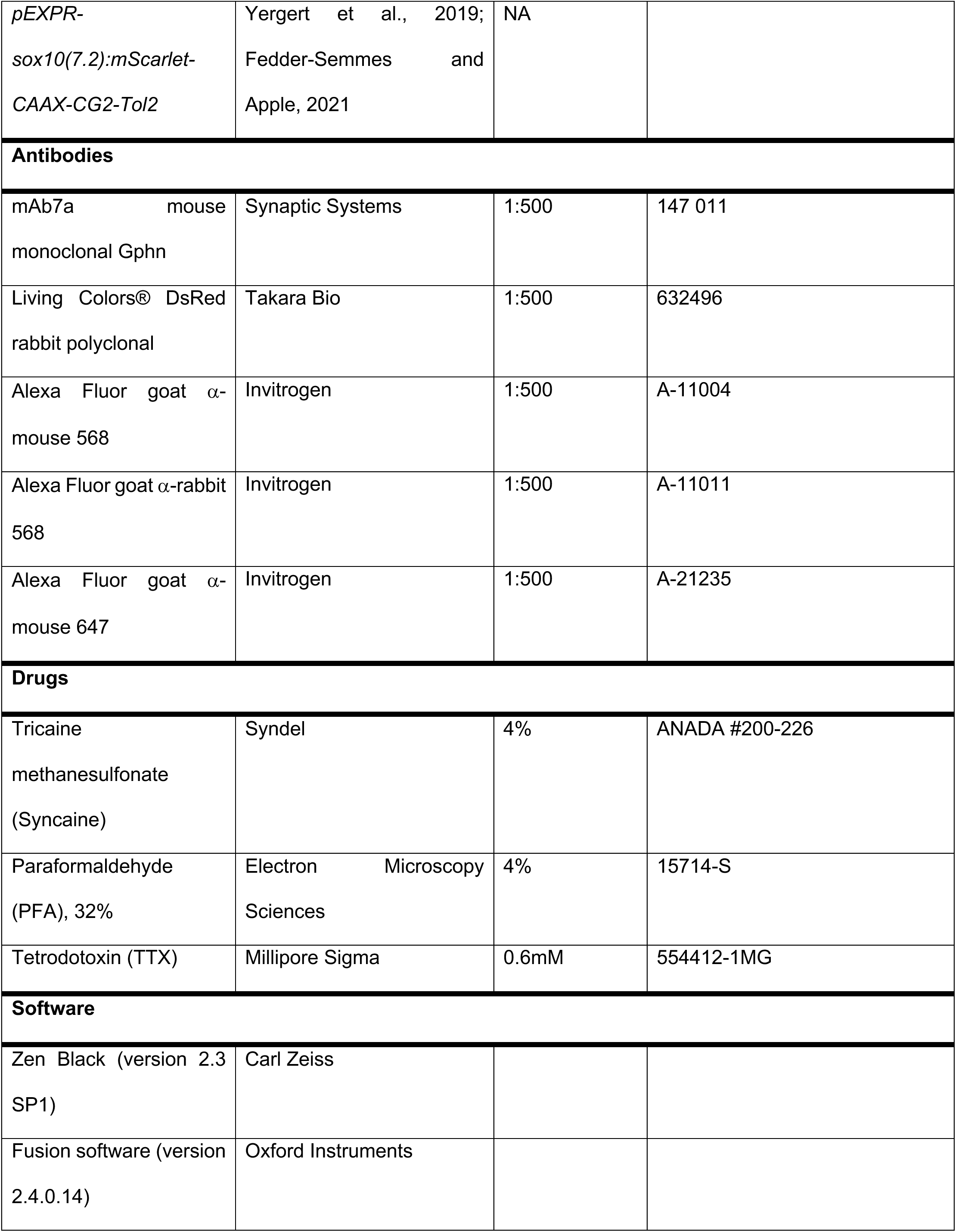
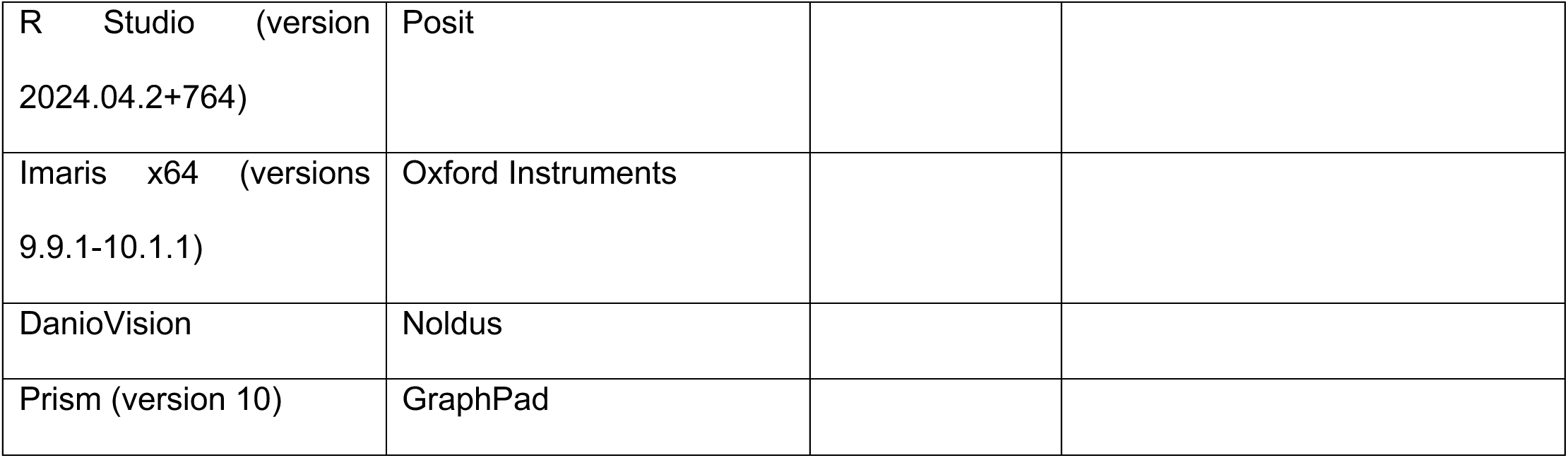

## Acknowledgments

We thank the Appel lab for comments on the manuscript. We also thank Dr. Angie Ribera for the *Tg(slc6a5:eGFP)* line, Dr. Rolf Karlstrom for the *TgBAC(gad1b:eGFP)* line, Dr. Don Arnold for *pCAG_GPHN.FingR-mKate2-IL2RGTC*, Dr. Thomas Schilling for *huβB1cry:mApple-CAAX*, and Dr. Anne Brunet for *pECE-M2-GPHN*. Finally, we thank the superb staff in our Zebrafish Core Facility for animal care.

## Author contributions

N.J.C. and B.A. conceived the project. C.A.D performed behavior experiments and analysis. B.A. made the *pME-mApple-CAAX* and *p3E-T2A-mApple-CAAX* plasmids. N.J.C. designed and performed all other experiments and analyzed all other data. N.J.C. wrote, and N.J.C., C.A.D., and B.A. edited the manuscript.

## Data, code, and materials availability

All data are available in the manuscript, Extended Data, or Source Data files. Raw images, R code, plasmid constructs, and zebrafish lines are available upon request.

